# Control of two-pathway signal integration in a model neocortical pyramidal cell

**DOI:** 10.1101/2025.04.21.649759

**Authors:** Bruce P. Graham, Jim W. Kay, William A. Phillips

## Abstract

We demonstrate how neuromodulation and spatially targetted inhibition can alter the integration of the two streams of excitatory input in a thick-tufted layer 5 pyramidal cell, using a computational reduced-compartmental cell models. Choosing suitable ranges of brief current stimulus amplitudes, applied basally to either the soma or basal dendrites (basal stimulation) and to the apical tuft (apical stimulation) results in burst firing due to either stimulus alone, if strong enough, or by a combination of the stimuli at lower amplitudes. Applying tonic inhibition to the apical tuft removes the ability of apical input alone to generate a burst over the chosen amplitude range. A similar effect is achieved by reducing the tuft calcium channel conductance as an outcome of neuromodulation. Similarly, tonic inhibition to the basal dendrites removes the ability of basal stimulation alone to generate a burst, without blocking bursts resulting from apical calcium spikes. HCN channels in the apical dendrites may amplify or reduce bursting probability, depending on other active and passive properties of dendrites. So neuromodulation that decreases the conductance of these channels may act to reduce or increase bursting probability across the across the ranges of basal and apical inputs, depending on cell properties. These effects mimic those found previously by simply limiting the range of stimulus amplitudes (***Graham et al., 2025***) but now show that such changes in two-stream signal integration can happen through network inhibition and neuromodulation with no change in the excitatory driving stimulus strengths. These changes in signal integration also lead to changes in information transmitted by the cell’s bursting probability about the two input streams, as shown in ***Graham et al. (2025)***. Changes in cell morphology are also investigated by reducing the apical trunk length and are revealed to alter this two-stream signal integration through differential effects on passive and active interaction between the soma and apical tuft.

## Introduction

In ***Graham et al. (2025)*** we used computer simulations of a reduced-compartmental model of a thick-tufted layer 5 neocortical pyramidal cell (***Bahl et al., 2012***) to investigate how the two major excitatory input streams, to the basal dendrites and apical tuft dendrites, respectively, interact to generate burst firing output. In essence, a short, high frequency burst of output spikes can arise through sufficiently long and strong basal input alone; or by strong apical input causing a dendritic calcium spike that propagates passively to the soma and there generates a burst of sodium spikes; or by back-propagation-active calcium spike firing (BAC-firing) where basal input generates a single somatic spike that back-propagates into the apical dendrites and combines with moderate apical input to generate a calcium spike, which then leads to an output burst (***Larkum, 2013***).

On these functional grounds for generating bursts, we defined four bursting regimes on the basis of the ranges of strength of the basal and apical inputs, for brief, synchronous input on both streams (***Graham et al., 2025***). High (H) ranges of input were defined up to amplitudes (strengths) that were sufficient for one input stream alone to induce a bursting output, thus giving an all-encompassing HH bursting regime in which input streams either alone or in combination could generate bursting. Subsumed within this, we defined low (L) ranges of input up to amplitudes just below that required to generate a finite probability of bursting due to that input alone. This gives three further regimes: HL in which basal inputs cover the full range (H) and can generate bursting alone, but low apical inputs (L) cannot; LH is vice versa with apical inputs (H) able to generate bursting alone but basal inputs (L) cannot; and LL in which bursts are only generate by basal and apical inputs in combination.

These regimes could each be described by transfer functions with distinct characteristics: LL is a combined function of basal and apical input; HL is a function of basal input only summed with a combined function of basal and apical input; LH is vice-versa, a function of apical input only summed with a combined function of basal and apical input; and HH is the sum of a function of basal input, a function of apical input only and a combined function of basal and apical input (***Graham et al., 2025***). This shows that HL and LH, in particular, are modulatory regimes in which the effect of the stronger input is modulated by the weaker, with the weaker input alone not able to generate a significant output.

Partial information decomposition (***Williams and Beer, 2010***) of bursting probabilities also reveals different information transmission characteristics within these four regimes (***Graham et al., 2025***). In the HH and LL regimes, bursting probability carries significant shared mechanistic and synergistic information about the strengths of the basal and apical input streams; whereas the HL and LH regimes transmit significant unique information about the stronger basal (HL) or apical (LH) input stream, with little or no unique information about the other stream.

### Inhibition and neuromodulation

Only excitatory basal and apical input streams were used in ***Graham et al. (2025)***. We now assume that these inputs may always cover the full HH ranges and we demonstrate how neuromodulation and spatially-targetted inhibition can alter the bursting regime due to these two streams of input in model layer 5 pyramidal cells. In particular, we explore how the subregimes HL, LH and LL can arise through the action of network inhibition and neuromodulation of membrane-bound ion channels, without explicitly restricting the ranges of the excitatory input amplitudes. These regimes can then be explicitly identified with particular behavioural states in wakefulness and sleep.

In neocortical layer 5 and other pyramidal cells, the interaction of apical and basal excitatory inputs is strictly controlled through inhibition, mediated by a variety of classes of interneuron that make layer-specific connections onto a pyramidal cell and are preferentially driven by top-down, bottom-up or lateral connections (***Gentet, 2012***; ***Kubota et al., 2016***). In particular, the apical tuft and perisomatic regions are subject to distinct spatially-targetted inhibition that limits the summation of excitatory inputs to those regions.

Pyramidal cells are also affected by a variety of neuromodulators whose concentration may vary with behavioural state on long (hours) and short time scales (seconds or less) (***Aru et al., 2020***). These include acetylcholine, noradrenaline, seratonin and dopamine. Through second-messenger pathways these neuromodulators can affect the intrinsic membrane properties of pyramidal cells in a variety of ways, including the regulation of calcium and HCN channel activity. Here we study the effects of changes in calcium and HCN channels on the generation of regenerative calcium spikes in the apical dendrites and the interaction of apical and basal excitatory inputs. Such interaction is influenced by HCN channels present with increasing density along the apical dendrites from the soma to the apical tuft.

In the following, we explore the effects of targetted inhibition and regulation of calcium and HCN channels on the burst firing probability due to brief excitatory basal and apical inputs.

### Apical trunk length

Apical trunk length has been shown both experimentally and in computational modelling to strongly influence the interaction between basal and apical inputs, particularly through the ability or otherwise to generate BAC-firing (***Fletcher and Williams, 2019***; ***Galloni et al., 2020***). Smaller layer 5 pyramidal cells tend not to burst at all but apical inputs can generate somatic single spike firing (***Fletcher and Williams, 2019***). Electrical compactness in these cells is enhanced by a reduced intracellular resistance along the trunk (***Fletcher and Williams, 2019***). It is unclear whether there are differences in calcium channel expression in the tuft (***Fletcher and Williams (2019)*** do not explore calcium influx directly), but it is possible that the increased electrical compactness means any calcium current leaks from a hotspot too quickly to generate a regenerative calcium spike. This is born out by the results of ***Galloni et al. (2020)*** who explore the effects of trunk length in both detailed models (large thick-tufted layer 5 ***Hay et al. (2011)*** model and the same biophysics applied to a reconstructed smaller PC) and in the ***Bahl et al. (2012)*** reduced model. The small detailed cell model does not generate a regenerative calcium spike in the apical tuft for any amplitude of apical input, and the resulting voltage transient does not cause even a single somatic spike, in contrast to the experimental results of ***Fletcher and Williams (2019)***. The ***Bahl et al. (2012)*** reduced model with a simple reduction in trunk length continues to produce a full calcium spike in the tuft, albeit with an increased generating rheobase current, which then causes output burst firing. ***Galloni et al. (2020)*** show that BAC-firing is actually more likely in larger (longer) cells due to the active back-propagation of a somatic sodium spike resulting in a broader spike at the apical tuft that is more effective in combining with apical excitation and generating a dendritic calcium spike.

Here we further explore the effect of trunk length in the bursting ***Bahl et al. (2012)*** reduced model. The results highlight the importance of the balance between active and passive interactions between apical and basal inputs in generating burst firing.

## Methods

This work uses the same methods as described in detail in ***Graham et al. (2025)*** (see also a preprint version stored on bioRxiv as ***Graham et al. (2024)***). A summary of these methods is given here.

### Computational cell models

Computer simulations of the ***Bahl et al. (2012)*** reduced-compartmental model and the ***Hay et al. (2011)*** detailed compartmental model of a thick-tufted layer 5 neocortical pyramidal cell are used to explore the contributions of basal and apical input streams to spiking output generation and bursting. The code bases for both models were obtained from ModelDB (https://modeldb.science). Most results in this work concern the ***Bahl et al. (2012)*** model.

The model cells were made noisy by insertion of two point current sources, one in the cell body and the other in the apical tuft, with the currents described by an Ornstein-Uhlenbeck (OU) process, as used by ***Larkum et al. (2004)***. Both noise sources had a mean current of 0 nA, with a standard deviation of 0.1 nA and a correlation length of 3 ms. The code was adapted from ***Destexhe et al. (2001)***.

In certain simulations, the model for the *I_h_* current used in the ***Bahl et al. (2012)*** model was augmented with a tonic leak current *I_lk_* whose conductance was proportional to the maximum HCN conductance, as per ***Migliore and Migliore (2012)***, giving the current: *I_lk_* = *c_lk_g-_h_*(*x*)(*V* − *E_lk_*), where *x* is the spatial position in the cell.

### Inputs

Excitatory inputs were given as simultaneous, brief point current injections to the soma or basal dendrites and the apical tuft or its base, over different ranges of amplitude. The form of current injection at each site is that used in BAC-firing experiments (***Larkum, 2013***) and consists of a square current pulse (duration 10 ms) to the soma or basal dendrite and an EPSC-like dual exponential waveform (rise time 0.5 ms; decay time 5 ms) to the apical dendrite.

Stimulus amplitude ranges for the ***Bahl et al. (2012)*** cell model cover the HH operating regime when no modulation of the cell is included: basal input from 0 to 1 nA and apical input from 0 to 1.7 nA. Every possible pair of input amplitudes within these ranges was trialled, with a basal increment of 0.05 nA and an apical increment of 0.1 nA, giving 21 × 18 data points.

For the ***Hay et al. (2011)*** cell model, the apical stimulation is given either at the distal end of the apical trunk (compartment 36), within the calcium hotspot in the apical tuft (compartment 60) or distal to the hotspot (compartment 74). In each case the apical noise source is inserted in the same compartment as the excitatory stimulus. Stimulus amplitude ranges again cover the HH operating regime when no modulation of the cell is included: basal input from 0 to 2 nA and apical input from 0 to 2 nA, in increments of 0.2 nA for both, giving 11 × 11 data points.

### Outputs

Simulations were run either 100 times (***Bahl et al. (2012)*** model) or 50 times (***Hay et al. (2011)*** model) for each pair of input current amplitudes but with different noise each time. Output spikes following the occurrence of excitatory input stimuli were counted individually and as a burst if two or more spikes occurred with an interspike interval of less than 25 ms.

Burst probabilities were calculated from the number of detected output bursts over the repeated simulations for each combination of amplitudes of basal and apical input for discrete increments in their respective amplitudes within their defined ranges.

### Transfer functions

Transfer functions for burst probability were defined in ***Graham et al. (2025)*** and these functions are used to here to fit to burst probability landscapes as appropriate. Fitting was carried out using Scipy’s least_squares method (scipy.org) in Python. Details of how these functions were derived are given in ***Graham et al. (2025)*** and the functions themselves are summarised here in the Appendix.

### Partial information decomposition (PID)

The information transmitted by the bursting output about the two input streams was quantified using partial information decomposition (PID), first formulated by ***Williams and Beer (2010)***. PID attempts to augment the standard Shannon mutual informations in a multivariate system by estimating information contained in the output that is unique to each input, shared by the inputs or is synergistic (only present due to the confluence of the inputs). Further details of this approach and the computational techniques used to evaluate a PID are given in ***Graham et al. (2025)***.

Results presented here were generated by the Ibroja decomposition method (***Bertschinger et al., 2014***; ***Griith and Koch, 2014***), which was implemented using compute UI (***Banerjee et al., 2018***) and the discrete information theory library dit (***James et al., 2018***). Python code was called from RStudio (R Core Team et al., 2021) by using the reticulate package (***Ushey et al., 2020***). The graphics were produced by using the ggplot2 package (***Wickham, 2016***) in RStudio.

## Results

### Original model

Figure 1 shows the burst probabilities estimated from simulation data from the unmodified ***Bahl et al. (2012)*** model, firstly as coloured contours (Figure 1a,b) and then as a function of basal amplitude for slices of apical amplitude (Figure 1c) and vice-versa (Figure 1d). Dotted blue lines in Figure 1a show the chosen boundaries between the low (L) and high (H) ranges for apical and basal input, which are also shown in solid blue in Figure 1c,d.

**Figure 1.**
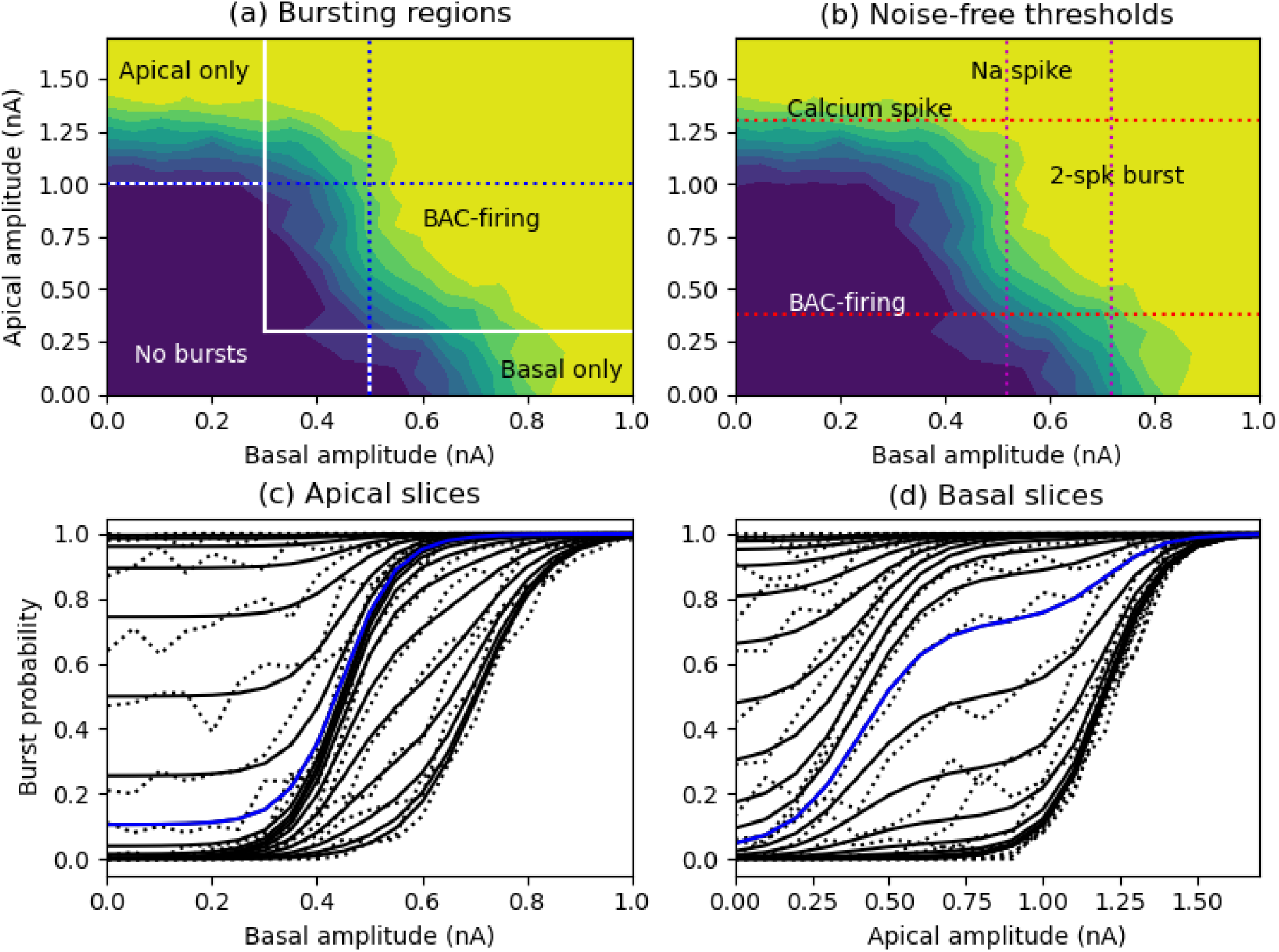
(a) Contour plots of burst probability from the simulation data for the HH operating regime. White lines: different bursting regions; blue dotted lines: qualitative boundaries between low (L) and high (H) input amplitudes (horizontal line: apical boundary; vertical line: basal boundary). (b) Contour plots of burst probability now overlayed with spiking thresholds in the noise-free cell (red dotted lines). Na spike: basal stimulation threshold for sodium spike in soma; Calcium spike: apical stimulation threshold for calcium spike in apical tuft; BAC-firing: apical stimulation threshold for calcium spike and subsequent somatic spikes following basal initiation of a first somatic spike; 2-spk burst: basal threshold for a somatic two-spike burst. (c) Simulation (dotted lines) and fitted transfer function (solid lines) curves for burst probability as a function of basal amplitude for individual values of apical amplitude from 0 to 1.7 nA in increments of 0.1 nA, bottom to top. (d) As for (c) but as a function of apical amplitude for values of basal amplitude from 0 to 1.0 nA in increments of 0.05 nA, bottom to top. Blue lines indicate selected boundaries between (c) HL and (d) LH regimes.

There are essentially three phases of burst firing evident in this burst probability landscape (Figure 1a, white lines): (1) Apical-only bursting due to apical input alone causing a dendritic calcium spike, which can happen for apical amplitudes around 1.0 nA and above; (2) Basal-only bursting due to basal input alone causing a rapid two-spike burst, which can happen for basal amplitudes around 0.6 nA and above; (3) BAC-firing bursting, which requires both basal and apical amplitudes of sufficient amplitude, with basal input around 0.3 nA and above generating an initial somatic spike which back-propagates and summates with sub-calcium-spike threshold apical input of around 0.3 nA and above). The associated sodium and calcium spike thresholds to basal and apical inputs, respectively, in the noise-free cell are shown in Figure 1b, to further indicate how these bursting regions emerge (see Table 1 in the Appendix for quantitative values of these thresholds).

**Table 1.** Data on soma-tuft interaction in a noise-free cell with an apical trunk length of either 500 or 200 *µ*m.

| Situation | 500 $\mu\text{m}$ trunk length | 200 $\mu\text{m}$ trunk length |
| --- | --- | --- |
| Tuft 1 nA stimulus | tuft 30 mV; soma 4.8 mV | tuft 28.5 mV; soma 7.5 mV |
| Soma 0.4 nA stimulus | soma 12.2 mV; tuft 3.0 mV | soma 12.4 mV; tuft 5.7 mV |
| Soma single spike | tuft 24 mV | tuft 16 mV |
| without tuft calcium | tuft 24 mV | tuft 16 mV |
| without trunk sodium | tuft 8 mV | tuft 10 mV |
| Soma single spike threshold | soma 0.52 nA | soma 0.49 nA |
| with 0.5 nA tuft stimulus | soma 0.49 nA | soma 0.41 nA |
| with 0.8 nA tuft stimulus | soma 0.47 nA | soma 0.33 nA |
| Soma double spike threshold | soma 0.72 nA | soma 1.05 nA |
| BAC-firing tuft threshold | tuft 0.38 nA | tuft 0.78 nA |
| Calcium spike threshold | tuft 1.3 nA | tuft 1.3 nA |

To examine this in more detail, we firstly look at the apical slice plots (Figure 1c). For weak apical input (right-most curves), burst probability is largely a function of basal input producing a two-spike burst at high amplitudes. As apical amplitude increases, the probability of a single somatic spike due to basal input causing a burst via BAC-firing increases. This corresponds with the rise in burst probability at lower basal amplitudes (around 0.4 −0.6 nA) with modest apical input (up to around 0.6 nA). Burst probability is clearly a function of both basal and apical amplitudes in this region. Then there is a region, for apical amplitudes around 0.7 to 0.9 nA, in which burst probability is almost purely a function of basal amplitude, which determines the probability of an initial somatic spike. In this region, corresponding to the band of overlying slices in the middle of Figure 1c, apical amplitude is then sufficient to generate a burst via BAC-firing once an initial spike is fired, thus its exact amplitude is not strongly relevant. At higher apical amplitudes (1.0 nA and above), apical input alone may be sufficient to generate a large dendritic calcium spike and thus an output burst. Corresponding with this, burst probability becomes purely a function of apical amplitude once basal amplitude drops below the level at which it can generate an iniitial somatic spike (below around 0.3 nA).

These bursting phases are also evident in the basal slice plots (Figure 1d). At low basal amplitudes (no initial somatic spike) bursting is purely a function of apical input (right-most overlying traces in Figure 1c). As basal amplitude increases it causes a somatic spike leading to bursting due to BAC-firing for modest apical amplitudes (up to around 0.6 nA). The kink in the basal slices at around an apical amplitude of 0.6 nA shows the transitiion from BAC-firing to apical-only bursting. At larger basal amplitudes, basal input alone may be suient to generate a two-spike burst, or to trigger a BAC-fired burst.

To further highlight the different operating regimes, the burst probability plots for the HL, LH and LL subregimes are shown in Figure 2. These will serve for comparison with the effects of inhibition and ion channel modulation to be shown below.

**Figure 2.**
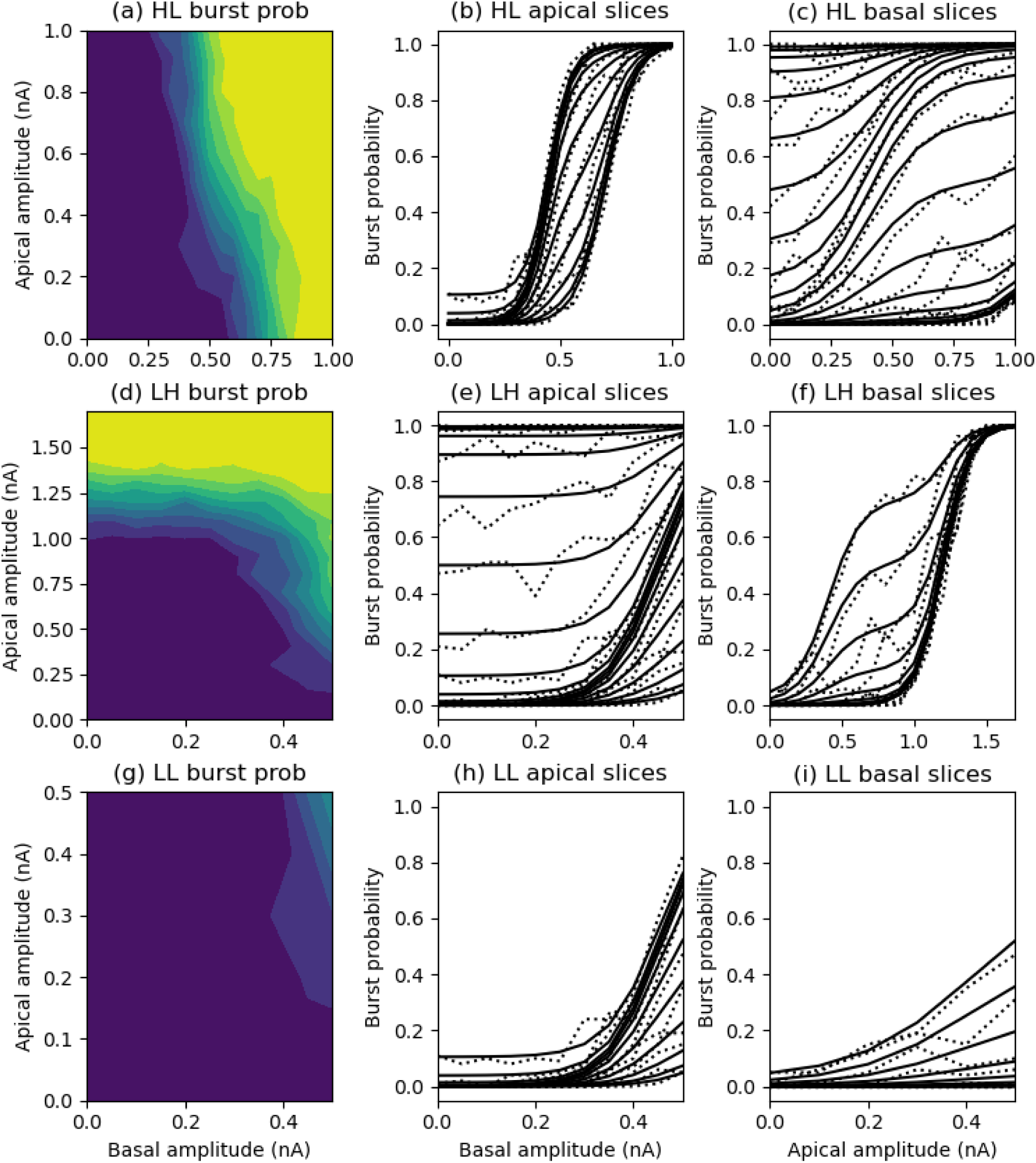
Contour and slice plots of burst probability from the simulation data restricted to the (a-c) HL region, (d-f) LH region and (g-i) LL region.

### Partial information decomposition (PID) analysis

PID covering the full ranges of basal and apical amplitudes for this original model is shown in Figure 3. Extending the Shannon mutual informations, PID provides estimates of the unique basal, unique apical, shared mechanistic and synergistic informations contained in the bursting output. Each point on the surface plots in Figure 3 indicates these PID quantities when the basal and apical amplitudes range from 0 up to the x-y value of the point.

**Figure 3.**
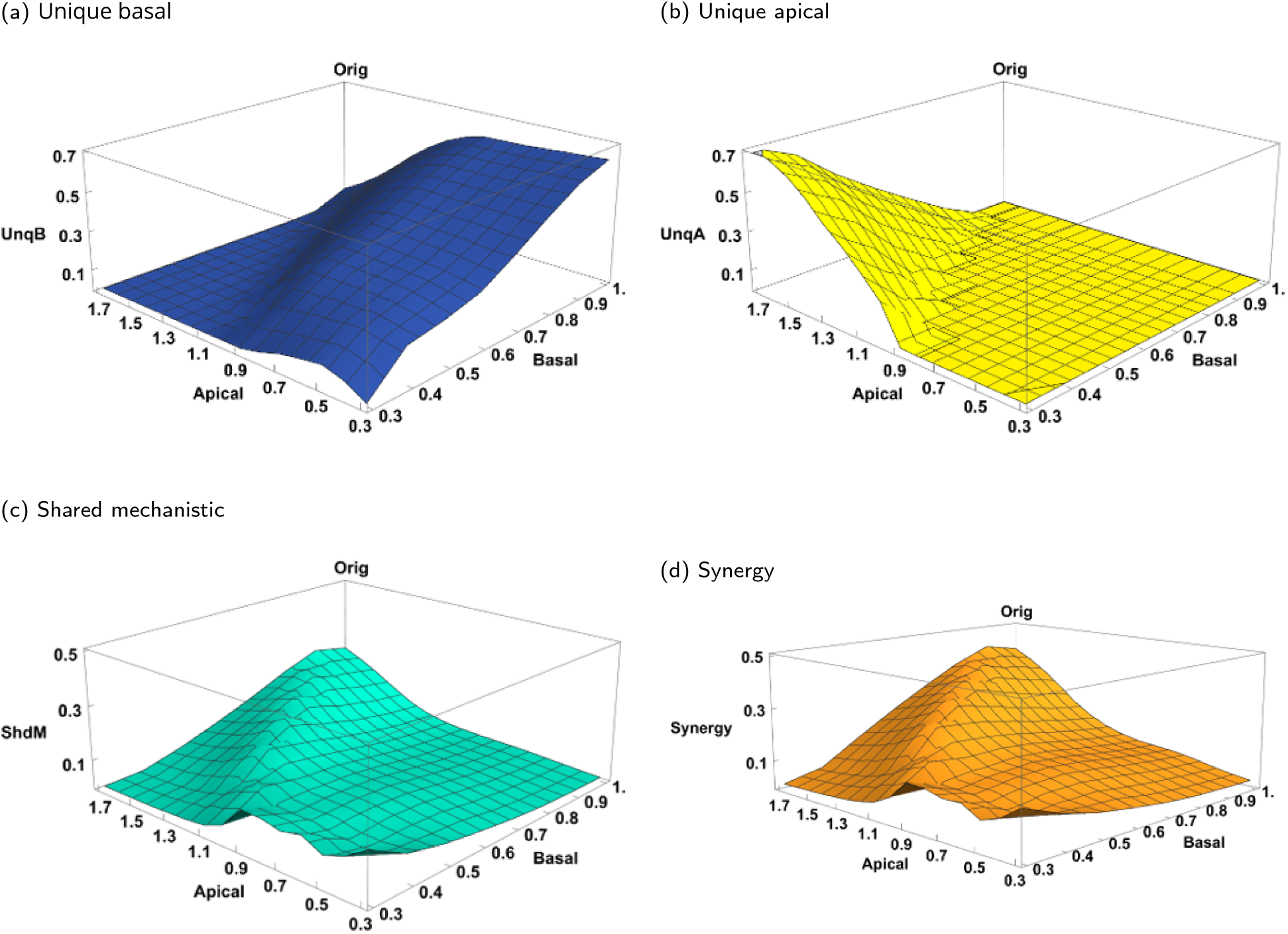
Normalised information-carrying PID quantities obtained by using the Ibroja method for pairs of basal and apical amplitude ranges: basal ranges from (0, 0.3), (0, 0.4) to (0, 1.0) nA; apical ranges from (0, 0.3), (0, 0.4) to (0, 1.7) nA, both in increments of 0.1 nA. (a) Unique basal. (b) Unique apical. (c) Shared mechanistic. (d) Synergistic.

It is apparent that there is an asymmetry between the information transmitted about the basal and apical inputs. No unique information about the apical input is transmitted in its low (L) range, below 1.0 nA. Rather, information transmission in the HL region (high basal above 0.5 nA and low apical amplitudes) is dominated by unique basal information. Then, in the LL region, with basal amplitudes below 0.5 nA, the is still unique basal information, but increasing shared mechanistic and synergistic information. This results because bursting is largely determined by the probability of a first somatic spike generated by basal input alone or by summed basal and apical input in their low ranges.

In the high apical range (amplitudes above 1.0 nA), information transmission largely contains unique apical information when basal amplitude is very low, as bursting output is almost purely determined by a dendritic calcium spike generated by the apical input alone. Unique apical information then declines and unique basal information begins to rise as basal amplitude increases and the BAC-firing regime is entered, which depends on a basally-triggered initial somatic spike. Also a clear ridge emerges along which shared mechanistic and synergistic information are the most significant, with unique information about either stream being low. This indicates that the apical and basal inputs along this ridge are making somewhat equal contributions to the occurrence of a burst, with basal input contributing at least the initial somatic spike and apical input responding to this with a dendritic calcium spike; or as both inputs become strong, either basal or apical input alone may generate the output burst, but it is not possible to tell which input path is chiefly responsible for any given burst.

In what follows we explore how inhibition and neuromodulation of membrane-bound ion channels can alter the operating regime of the cell in terms of the output burst probability and the consequent information transmitted about the two input streams.

### Inhibition

The results of tonic inhibition applied to the apical or basal dendrites, either singly or together, are shown in Figure 4. As is expected, such inhibition is able to limit the effective amplitude ranges of the excitatory basal and apical inputs, and thus shift the operating region from HH to HL, LH or LL, depending on the application of inhibition. More detail of these effects is given below.

**Figure 4.**
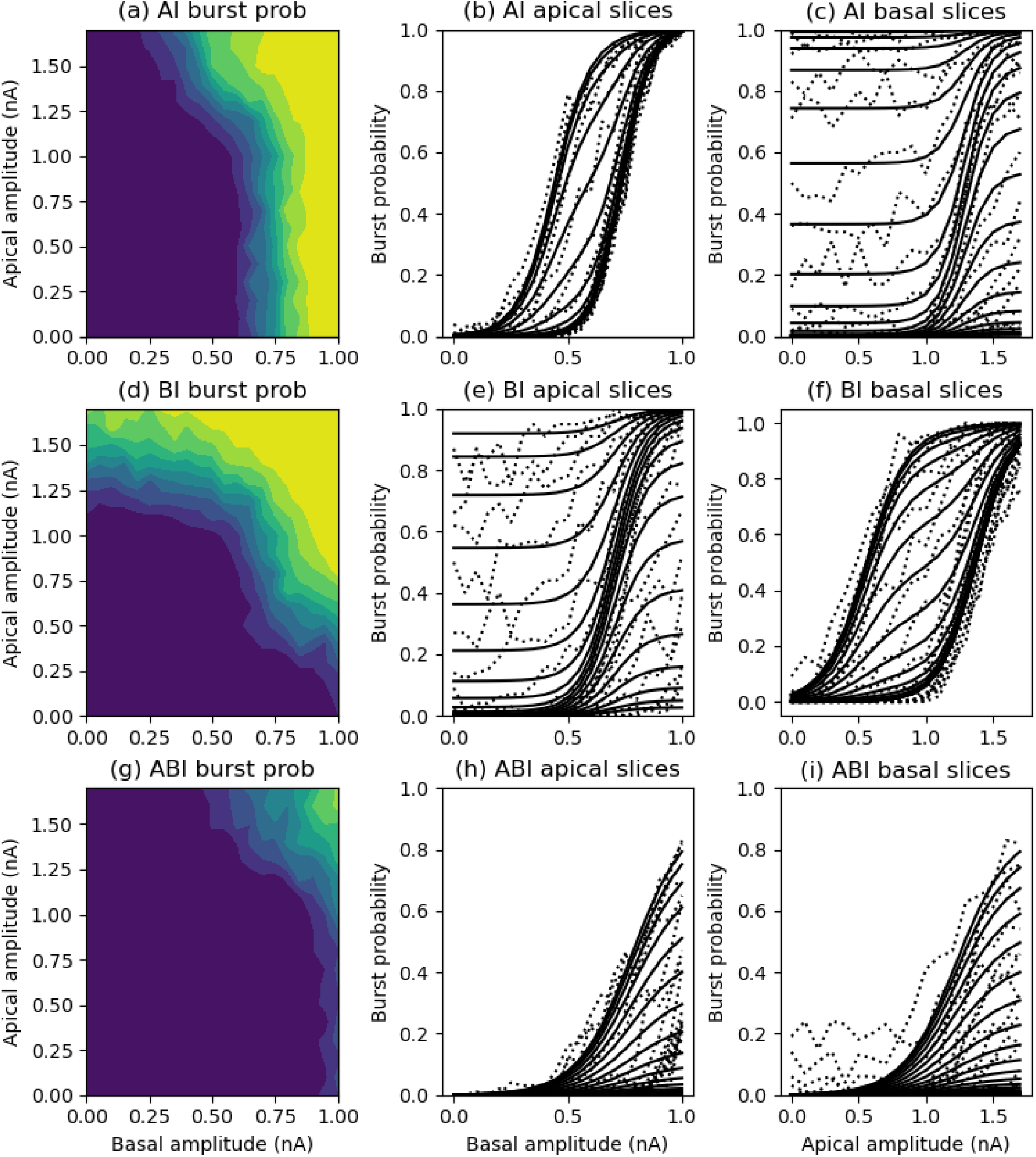
Contour and slice plots of burst probability from the simulation data for the HH amplitude ranges but with dendritic inhibition. (a) Tonic inhibition of 0.01 *µS* applied to the apical tuft. (b) Tonic inhibition of *µS* applied to the basal dendrites, with basal excitation shifted to the basal dendrites. (c) Tonic inhibition of a strength of 0.007 *µS* applied to both the basal dendrites and apical tuft. Solid lines in the slice plots are transfer function fits using the reduced transfer functions of the expected operating regimes (HL, LH and LL, respectively).

### Apical tuft inhibition

Addition of a tonic inhibitory conductance to the tuft, with a reversal potential of −75 mV, reduces the effect of apical input on cell excitability and thus produces a response that is very similar to the HL regime in the unmodified reference model. This is illustrated by the burst probability map and individual slice plots in Figure 4a-c. Values of apical amplitude up to 1.0 nA now have little effect on the burst probability due to basal input. Then in the apical range from 1.0 to 1.7 nA the apical input is strong enough to produce BAC-firing, despite the inhibition. This bursting landscape is comparable in form that of the HL regime (Figure 2a). The quality of fit of the HL transfer function (Figure 4b,c) is only a little worse than for the fitting of the full HH function (Appendix: Figure 14b,c and Table 2).

**Table 2.**
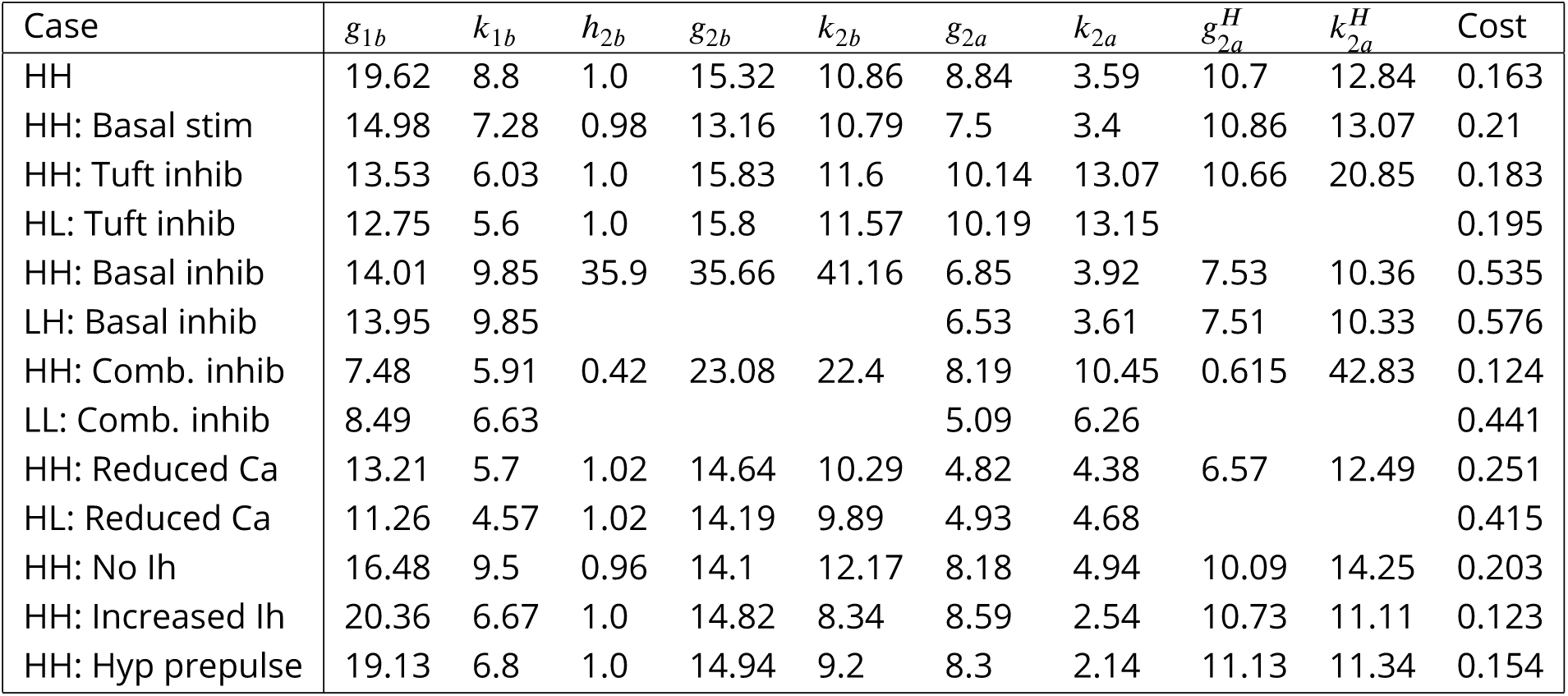
Least squares fits of the HH (*P*_2_(*b, a*)) transfer function parameter values for burst firing. Cost is the final sum-of-squares difference between the transfer function and the simulated data.

The PID analysis (Figure 5) shows largely the contextually-modulated processing profile of the HL regime (***Graham et al., 2025***), with significant unique information transmitted about the strength of basal input, with little unique information about the apical input, when basal input is above around 0.5 nA. When basal input is weak (below 0.5 nA), there is some unique information about the apical input when the apical amplitude ranges up to its highest amplitude (1.7 nA), but this much reduced compared with the uninhibited HH regime (Figure 3b). There is some significant shared mechanistic and synergistic information in this region (Figure 3c,d for apical amplitude up to around 1.0 nA). The similarity with the HL regime is further confirmed when comparing the PID breakdown over the whole amplitude ranges when apical inhibition is applied with that for the HL regime (apical and basal from 0 to 1 nA; see Appendix Figure 16).

**Figure 5.**
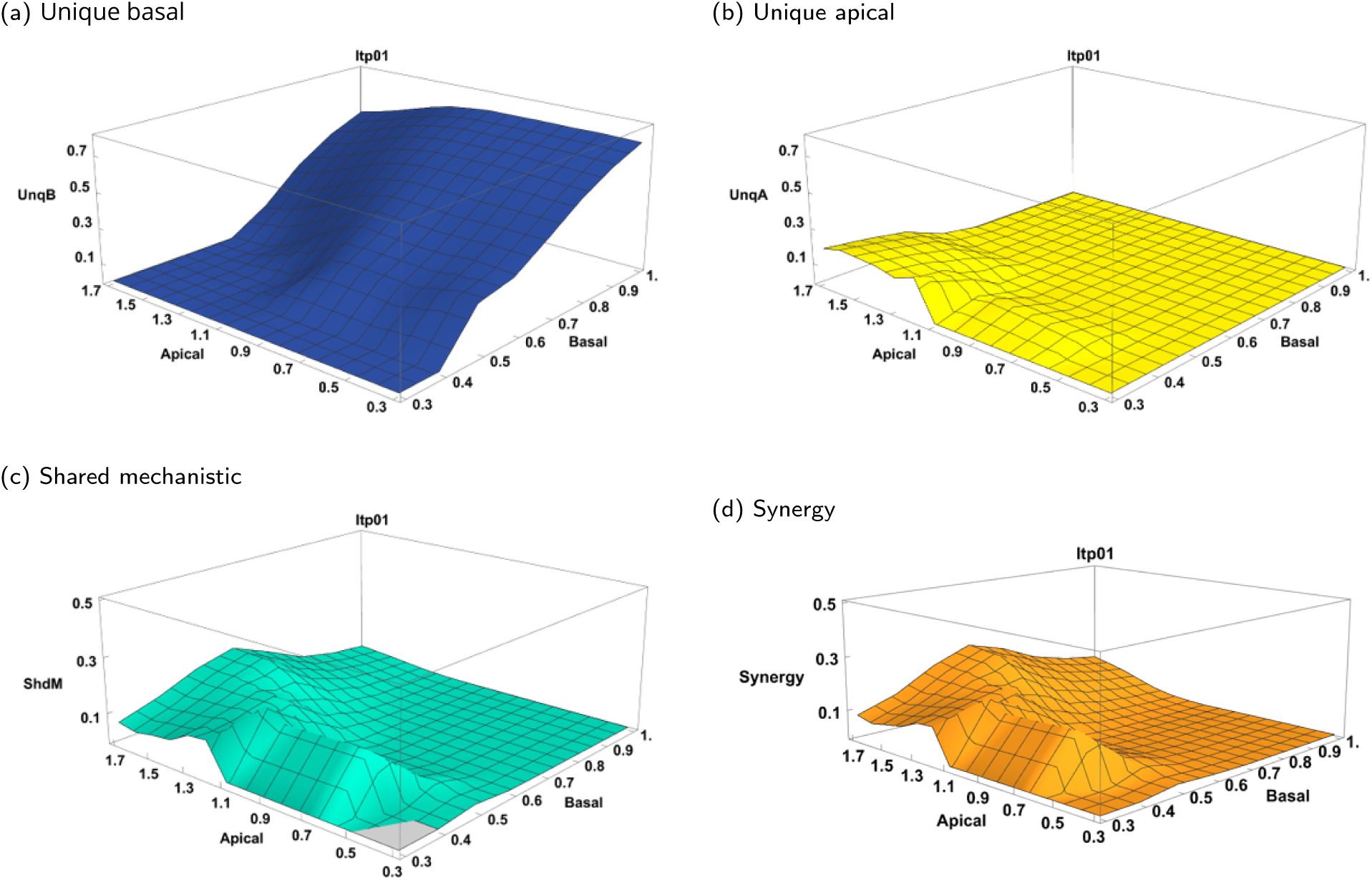
Normalised information-carrying PID quantities obtained by using the Ibroja method for pairs of basal and apical amplitude ranges when tonic inhibition of 0.01 *µS* is applied to the apical tuft. Basal ranges run from (0, 0.3), (0, 0.4) to (0, 1.0) nA; apical ranges run from (0, 0.3), (0, 0.4) to (0, 1.7) nA, both in increments of 0.1 nA. (a) Unique basal. (b) Unique apical. (c) Shared mechanistic. (d) Synergistic.

Thus this application of tonic inhibition to the apical tuft has reduced the bursting regime from one that one that allows bursting due to strong apical or basal input alone to one in which the burst probability due to basal input is contextually-modulated by the apical input, but without a change in the amplitude ranges of these excitatory inputs.

### Basal inhibition

To test the effect of perisomatic inhibition it is necessary firstly to move the basal current injection stimulation to the basal dendrites so that inhibition of the basal dendrites can impact basal excitatory input before its transmission to the cell body and subsequent interaction with apical input in the generation of cell output. Without inhibition, excitatory stimulation to the basal dendrites produces only a small quantitative difference in the output bursting profiles of the original somatic stimulation (not shown).

Addition of a tonic inhibitory conductance to the basal dendrites, with a reversal potential of −75 mV, reduces the effect of basal input on cell excitability and thus produces a response that is very similar to the LH regime in the unmodified reference model. This is illustrated by the burst probability map and individual slice plots in Figure 4d-f, which are comparable in form to those for the LH regime (Figure 2d-f). The ability of basal input alone to cause a burst is nearly eliminated, whereas strong apical input alone can produce a burst with high, though reduced probability.

Note that the fit of the LH transfer function (Figure 4e,f) is nearly as good as the fit of the full HH transfer function (Appendix: Figure 14e,f).

The PID analysis also shows a similar information processing profile to the LH regime (Figure 6; compare with Figure 3 for basal amplitudes up to 0.5 nA; for a point-wise comparison see Appendix Figure 16). The quantitative breakdown of PID component values shows less unique apical information is recovered via this strength of basal inhibition than is obtained in the LH regime (basal from 0 to 0.5 nA and apical from 0 to 1.7 nA; see Appendix Figure 16), but no attempt has been made to optimise the strength of inhibition with regards to this.

**Figure 6.**
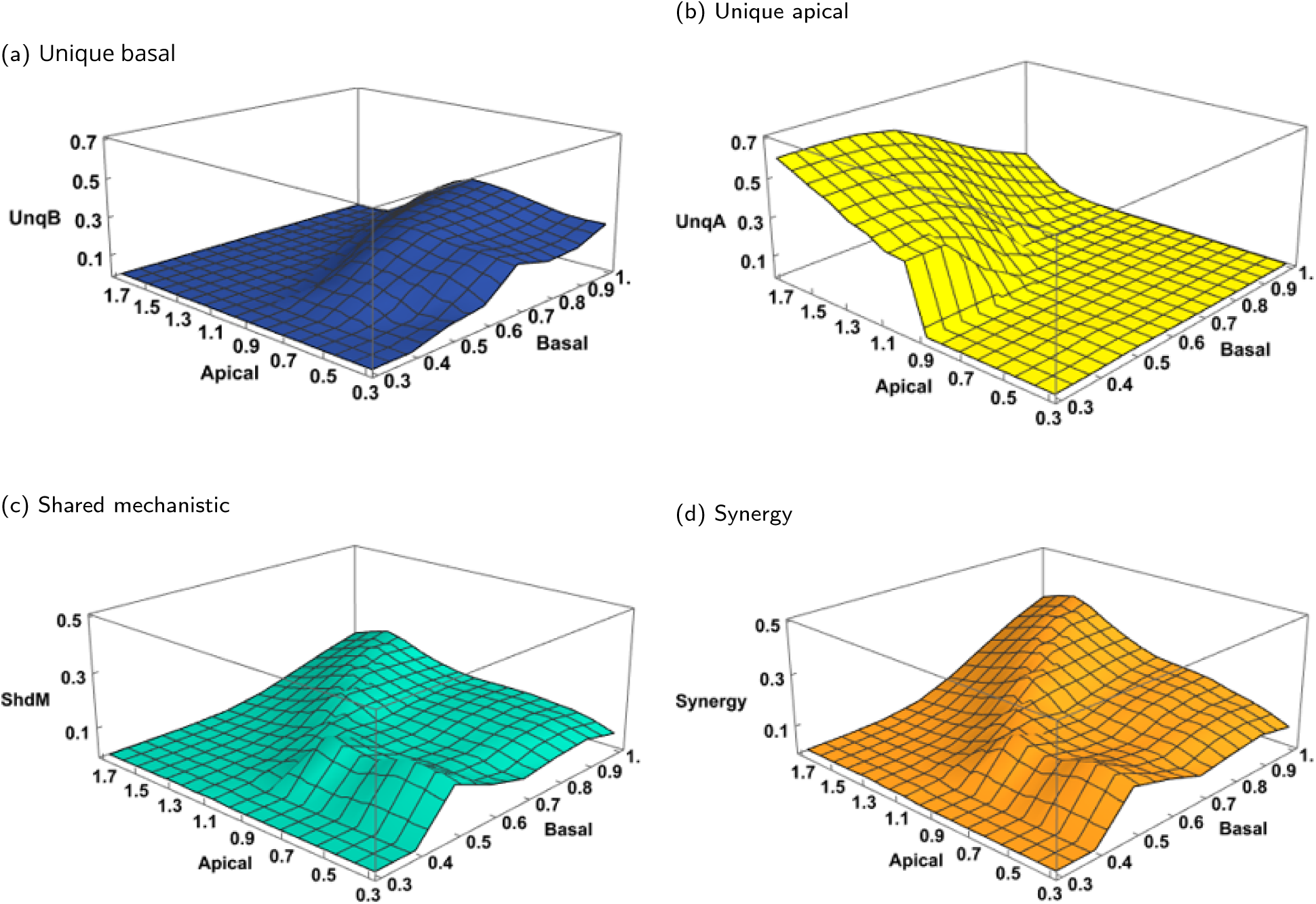
Normalised information-carrying PID quantities obtained by using the Ibroja method for pairs of basal and apical amplitude ranges when basal excitation and tonic inhibition of 0.01 *µS* is applied to the basal dendrites. Basal ranges run from (0, 0.3), (0, 0.4) to (0, 1.0) nA; apical ranges run from (0, 0.3), (0, 0.4) to (0, 1.7) nA, both in increments of 0.1 nA. (a) Unique basal. (b) Unique apical. (c) Shared mechanistic. (d) Synergistic.

Thus this application of tonic inhibition to the basal dendrites has reduced the bursting regime from one that one that allows bursting due to strong apical or basal input alone to one in which the burst probability due to apical input is modulated by the basal input, but without a change in the amplitude ranges of these excitatory inputs.

### Combined basal and apical inhibition

As expected, tonic inhibition to both the basal dendrites and apical tuft reduces burst firing to the LL range. Using a slightly reduced tonic inhibition amplitude (0.007 *µS*) in both basal and apical locations results in a bursting profile (Figure 4g-i) close to the original LL regime (Figure 2g-i). Note that the LL transfer function does not allow for basal or apical inputs alone resulting in a burst, but for these levels of inhibition the simulations show a small probability that basal amplitudes close to 1.0 nA can still cause bursting, so the fit of the LL transfer function (Figure 4i) is a little worse here than that of the full HH function (Appendix: Figure 14i).

The PID analysis also shows a similar information processing profile to the LL regime (Figure 7; compare with Figure 3 for basal amplitudes up to 0.5 nA and apical amplitudes up to 1.0 nA). A point-wise breakdown of PID values over the whole amplitude ranges is close to that of the LL regime (basal from 0 to 0.5 nA and apical from 0 to 1.0 nA; see Appendix Figure 16).

**Figure 7.**
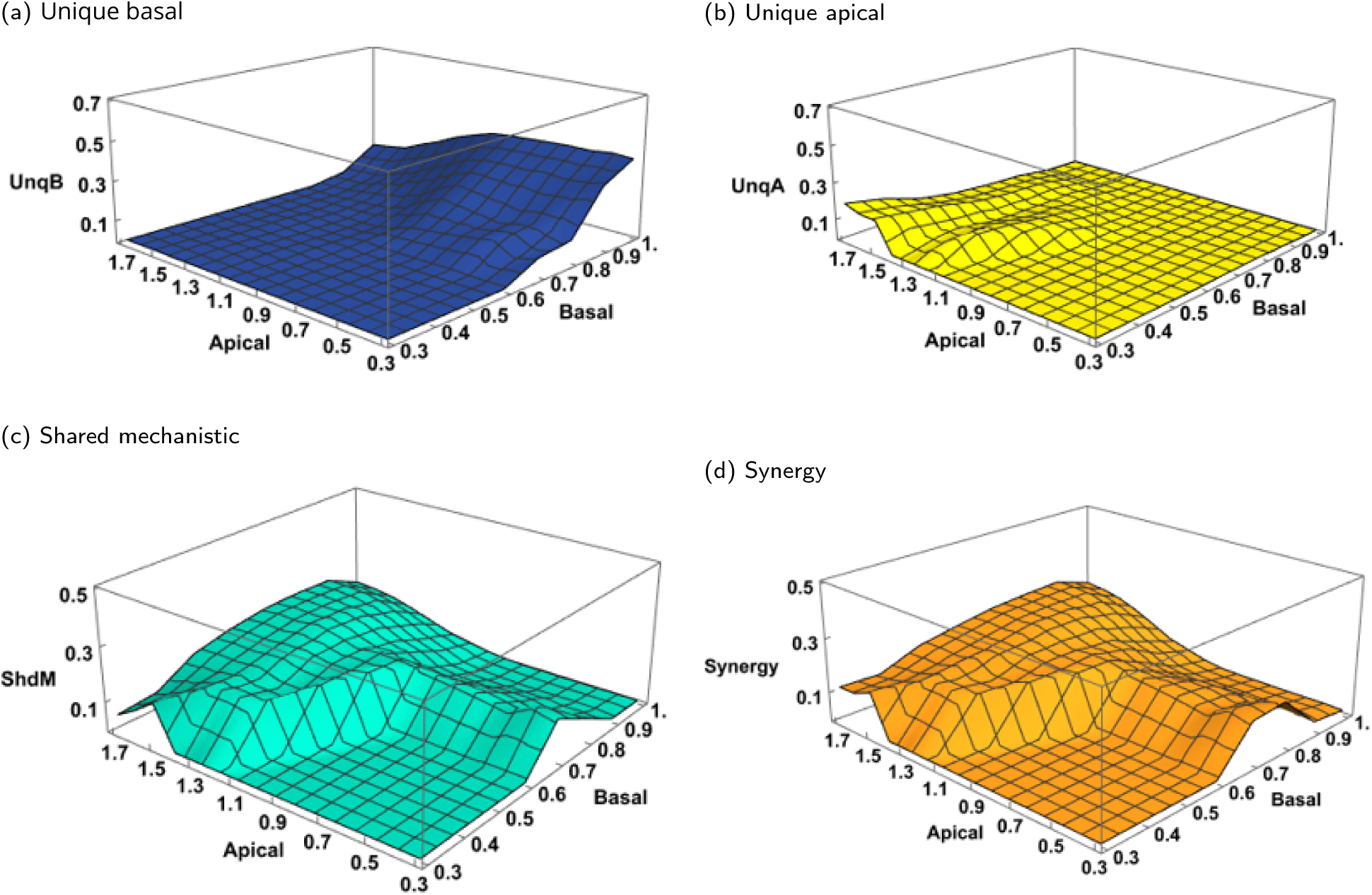
Normalised information-carrying PID quantities obtained by using the Ibroja method for pairs of basal and apical amplitude ranges when basal excitation is to the basal dendrites and tonic inhibition of 0.007 *µS* is applied both to the basal and apical dendrites. Basal ranges run from (0, 0.3), (0, 0.4) to (0, 1.0) nA; apical ranges run from (0, 0.3), (0, 0.4) to (0, 1.7) nA, both in increments of 0.1 nA. (a) Unique basal. (b) Unique apical. (c) Shared mechanistic. (d) Synergistic.

### Calcium channels

Reducing the density of calcium channels in the apical tuft, a form of down-regulation, results in smaller calcium transients for a given amplitude of apical input. As a consequence, such down-regulation of calcium channels in the apical tuft acts similarly to tonic apical inhibition. Reducing calcium channel density to a quarter of the original value effectively reduces the operating regime from HH to be more like HL (Figures 8. Dendritic calcium spikes may still be of sufficient amplitude to contribute to output spiking, thus allowing bursting through apical-only input and BAC-firing, but with reduced probability. Since the HL transfer function does not include the possibility of apical-only induced bursting, its fit to this data (Figure 8b,c) is slightly worse than that of the full HH transfer function (Appendix: Figure 15b,c).

**Figure 8.**
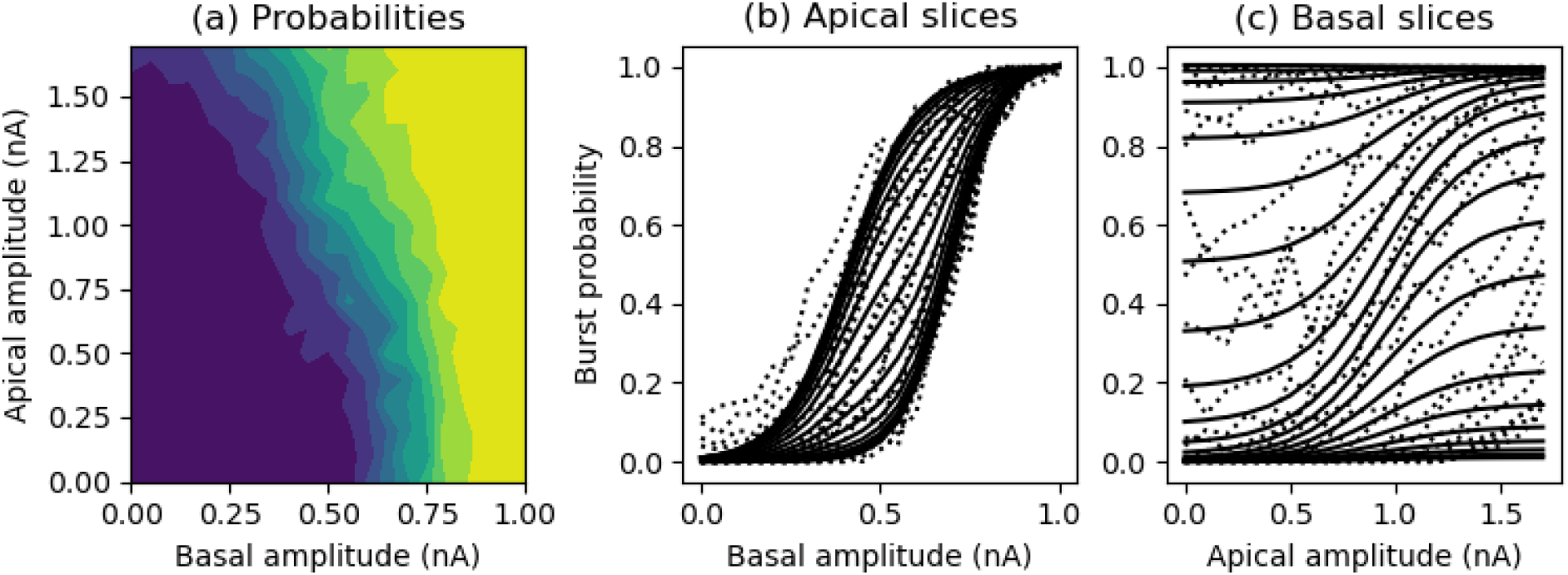
(a) Contour plots of burst probability from the simulation data for the HH amplitude ranges with tuft calcium channel density reduced to one quarter of its original value. (b) Simulation (dotted lines) and fitted HL transfer function (solid lines) curves for burst probability as a function of basal amplitude for individual values of apical amplitude. (c) As for (b) but as a function of apical amplitude for values of basal amplitude.

This reduction to the HL regime is born out by the PID profiles (Figure 9), which are very similar in form to those produced by apical inhibition (Figure 5). The similarity with the HL regime is further confirmed when comparing the PID breakdown over the whole amplitude ranges when calcium conductance is reduced with that for the HL regime (apical and basal from 0 to 1 nA; see Appendix Figure 16) and it is also similar to when apical inhibition is applied (Figure 16).

**Figure 9.**
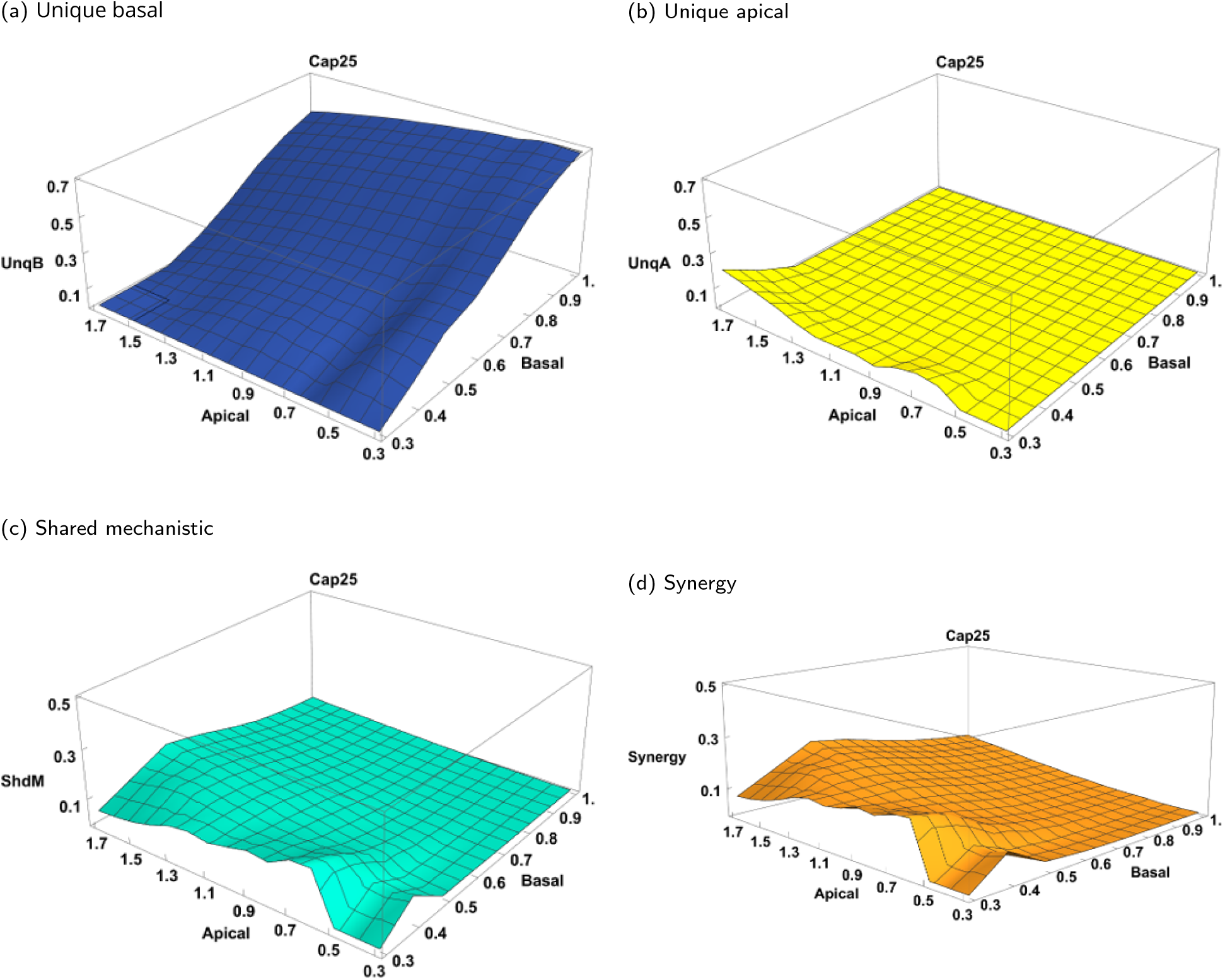
Normalised information-carrying PID quantities obtained by using the Ibroja method for pairs of basal and apical amplitude ranges when calcium channel density in the apical tuft is reduced to one quarter. Basal ranges run from (0, 0.3), (0, 0.4) to (0, 1.0) nA; apical ranges run from (0, 0.3), (0, 0.4) to (0, 1.7) nA, both in increments of 0.1 nA. (a) Unique basal. (b) Unique apical. (c) Shared mechanistic. (d) Synergistic.

This implies that if the cell operates in this state when cholinergic modulation is low, then it may enter the HH state when acetylcholine levels are high.

### HCN channels

In ***Bahl et al. (2012)*** model, HCN channel activity effectively increases cell excitability due to its depolarisation of the cell membrane, particularly in the apical tuft. The depolarising effect of these channels does not inactivate the high-voltage-activated (HVA) calcium channels and dominates over the decreased membrane resistance due to the H channel conductance, which could act to reduce excitability. Complete removal of the HCN channels decreases burst probability across both basal and apical axes (Figure 10b), thus approaching the LL regime (though the effect is not as strong as the effect of the reduced input amplitudes used in the reference model). Conversely, increasing the density of HCN channels boosts burst probability across both axes (Figure 10c), thus expanding the HH regime.

**Figure 10.**
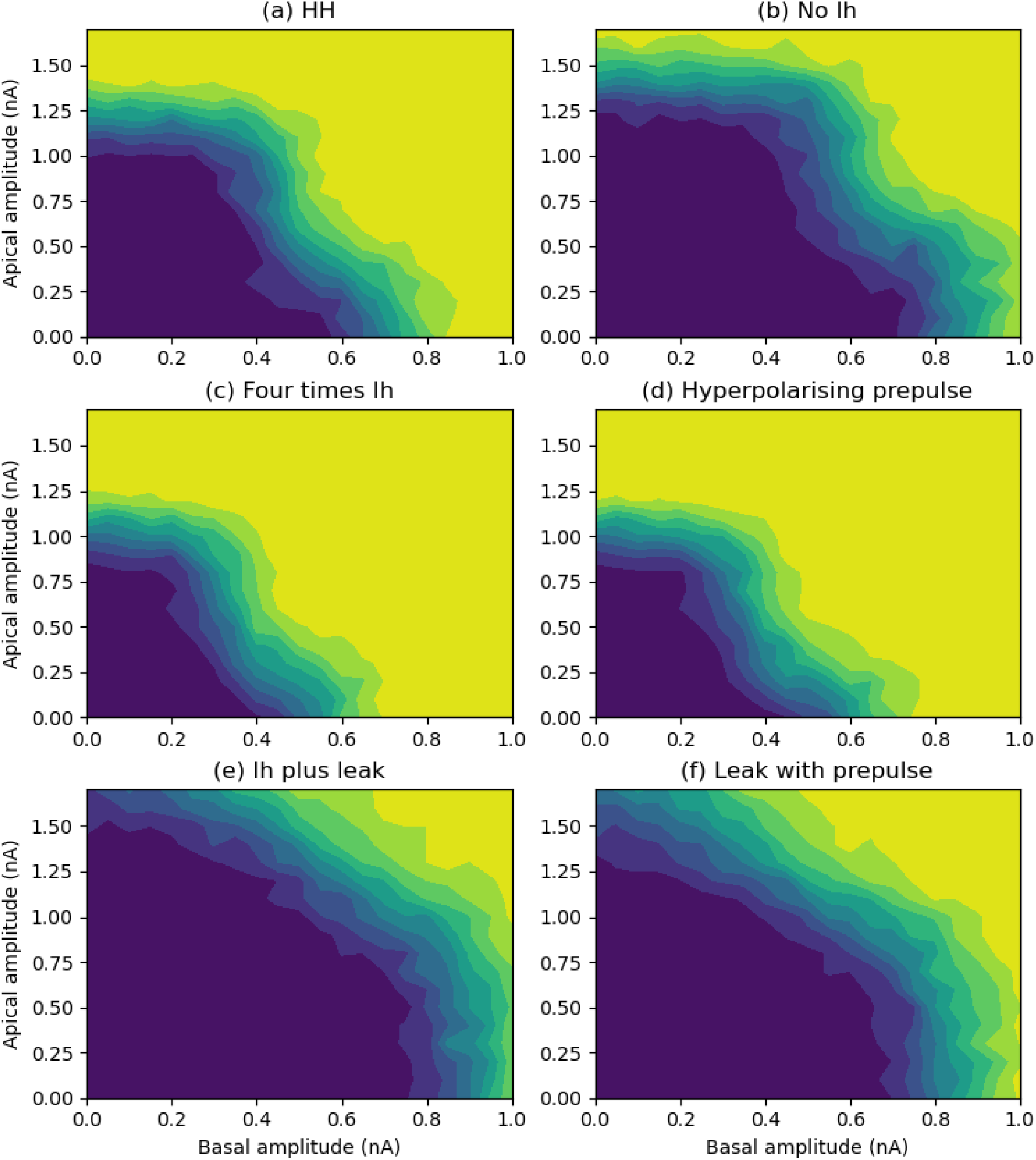
Contour plots of burst probability from the simulation data for (a) the HH operating regime; (b) with HCN channels removed; (c) with HCN channel density increased to four times its original value; (d) with original HCN channel density and with a 50 ms hyperpolarising current pulse of amplitude −1 nA that finishes 20 ms before the arrival of the basal and apical inputs; (e) inclusion of a tonic leak current with conductance proportional to the maximum HCN channel conductance (*c_lk_* = 0.03 and reversal potential *E_lk_* = −70 mV); (f) tonic leak current together with a 50 ms hyperpolarising current pulse.

This contrasts with experimental results that show that HCN channels have an inhibitory effect on apical dendrite stimulation, putatively due to the increased shunt provided by the HCN channel conductance (***Harnett et al., 2015***). Computational modelling work indicates that this shunt alone is insufficient (inline with our results for the ***Bahl et al. (2012)*** model) but that addition of a tonic leak current due to a conductance that is proportional to the maxiumum H channel conductance can provide sufficient shunt (***Migliore and Migliore, 2012***). Including such a leak current in the ***Bahl et al. (2012)*** model does indeed turn the effect of the HCN channels from excitatory to inhibitory for apical stimulation (Figure 10e).

These results apply only if the cell is in a (noisy) resting state when the excitatory stimuli are applied. The major characteristic of the HCN channels is that they fully activate at hyperpolarised levels and thus provide rebound excitation when signals driving hyperpolarisation cease. To test the effects of these HCN channel dynamics on bursting probability, trials were carried out where a hyperpolarising current injection was given to the apical tuft for 50 mS prior to the arrival of the excitatory stimuli. With the original reference density of HCN channels this results in a similar boosting of the burst probability as that obtained with an increase in HCN channel density (Figure 10d). An increase in burst probability is also seen when the tonic leak current is included, but not quite to the level obtained on removal of the HCN channels (Figure 10f).

### Detailed Hay cell model

It has been demonstrated by ***Mäki-Marttunen and Mäki-Marttunen (2022)***, using the ***Hay et al. (2011)*** detailed compartment model of a layer 5 pyramidal cell, that HCN channels can have either an excitatory or inhibitory effect, depending on the apical site of stimulation. Inhibition arises due to distal stimulation, where the high density of HCN channels depolarises the membrane at rest and so inactivates low-voltage-activated (LVA) calcium channels. Since these channels are not included in the ***Bahl et al. (2012)*** reduced model, to explore the possible inhibitory effects of HCN channels, we used the model of ***Hay et al. (2011)*** which includes low-voltage-activated (LVA) calcium channels in the apical dendrites and at high density in a calcium hotspot at the base of the apical tuft. The same stimulation protocols as used with the ***Bahl et al. (2012)*** model are applied, but with the apical site of stimulation trialled at different locations.

As with ***Mäki-Marttunen and Mäki-Marttunen (2022)***, the effect of the HCN channels depends on the location of the point stimulation to the apical dendrites, relative to the calcium hotspot (Figure 11). Comparing the top and middle rows of Figure 11, it can be seen that the amplifying effect of the HCN channels remains if apical stimulation is applied to the apical trunk just proximal to the calcium hotspot (apical model compartment 36). However, if apical stimulation is moved distally, either into the hotspot (model compartment 60) or distal to the hotspot (model compartment 74) then the density of HCN channels, which rises with distance from the soma, is sufficient to cause inhibition of calcium spike generation and subsequent somatic bursting, through inactivation of the LVA calcium channels.

**Figure 11.**
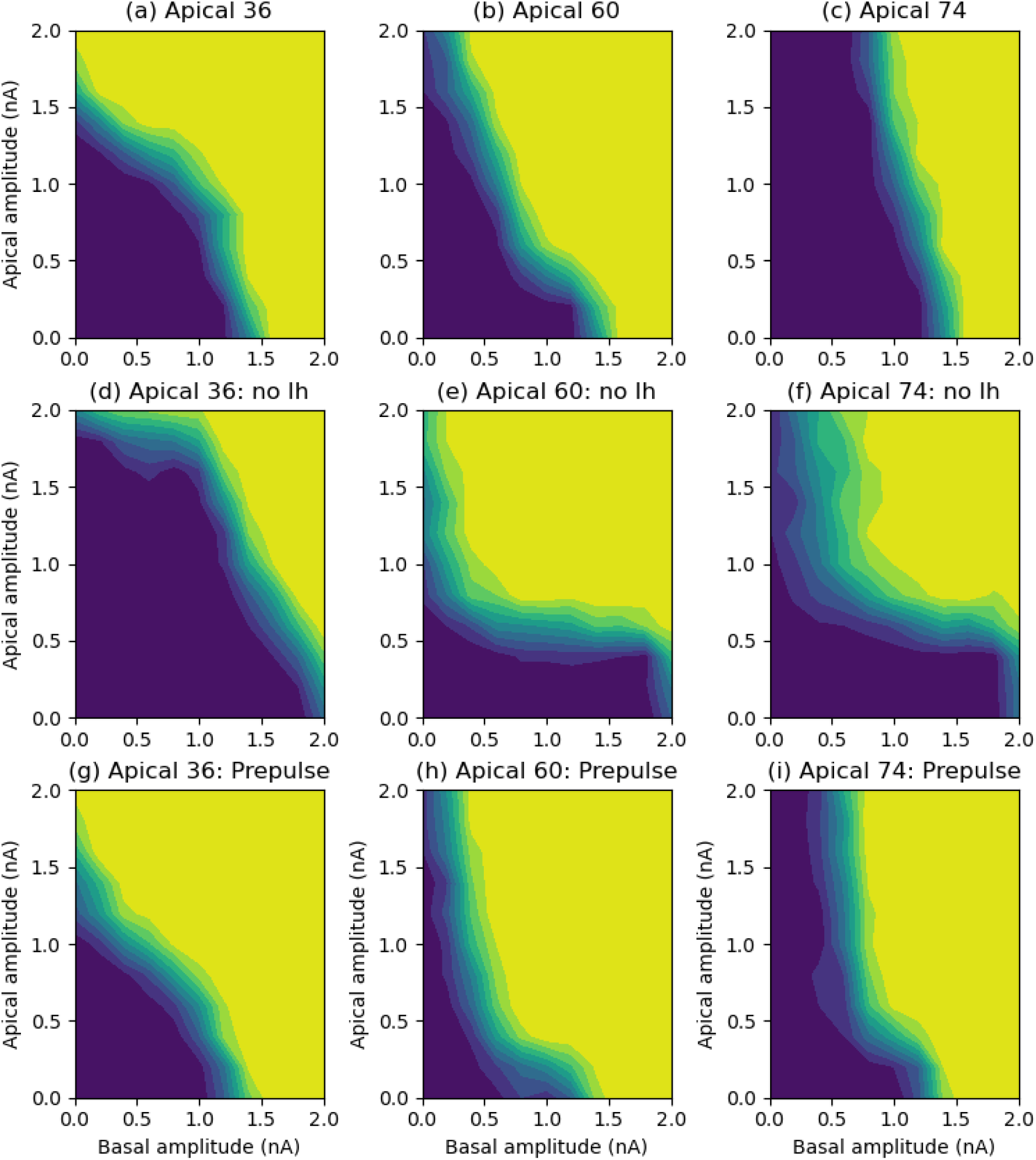
Contour plots of burst probability from the simulation data for the *Hay, et al.* (2011) cell model with apical stimulation at different points in the apical dendrites. (a) apical stimulation in model compartment 36 at 629 *µ*m from the soma, just proximal to the calcium hotspot; (b) apical stimulation in model compartment 60 at 817 *µ*m, inside the calcium hotspot; (c) apical stimulation in model compartment 74 at 1051 *µ*m, distal to the calcium hotspot; (d-f) as for (a-c) but with HCN channels removed; (g-i) with HCN channels, but with a 50 ms hyperpolarising current prepulse applied to the calcium hotspot (compartment 60) of amplitude −1 nA that finishes 20 ms before the arrival of the basal and apical inputs.

To test the effect of HCN channel dynamics, a 50 ms hyperpolarising current injection was given to the calcium hotspot (compartment 60) before the arrival of excitation, as trialled with the ***Bahl et al. (2012)*** model. As is evident in the bottom row of Figure 11, this prepulse now results in a boosting of burst probability for all three locations of apical stimulation. This is due to the activation of the HCN channels and the deinactivation of the LVA calcium channels by the hyperpolarisation. Thus the inhibitory effects of the HCN channels in the resting state are reversed.

### Short trunk length

The shortening the apical trunk length to 200*µ*m in the ***Bahl et al. (2012)*** model (from the original 500*µ*m), while preserving the biophysics, does not eliminate the ability of the cell to BAC-fire or to induce an apical calcium spike by apical input alone. However, the interactive effects between the soma and apical tuft are altered, leading to some counter-intuitive changes in the burst probability landscape across the range of basal and apical input amplitudes trialled. The comparison of this bursting landscape in the short and long cells is shown in Figure 12. These differences in passive and active soma-tuft interaction with trunk length result in a burst probability profile in the short cell that is a mirror image of that produced with a long trunk length, with the impacts of basal and apical inputs seemingly reversed. This is particularly apparent when comparing the slice plots for the short cell in Figure 12b,c, with those for the long cell in Figure 1c,d.

**Figure 12.**
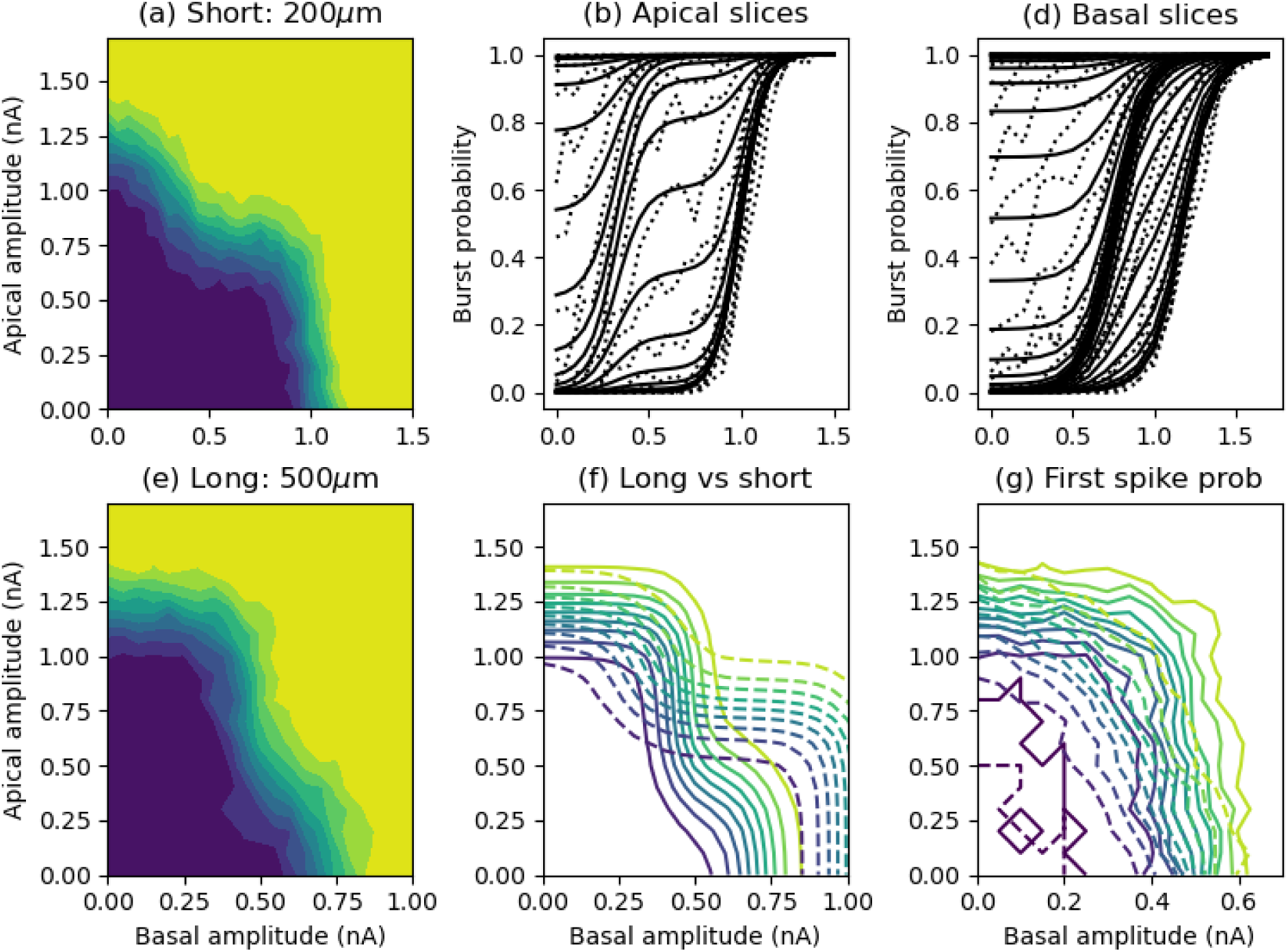
(a) Contour plots of burst probability from the simulation data for a short apical trunk length. (b) Simulation (dotted lines) and fitted transfer function (solid lines) curves for burst probability as a function of basal amplitude for individual values of apical amplitude. (c) As for (b) but as a function of apical amplitude for values of basal amplitude.

The difference between the two sets of contour plots is highlighted by the overlay in Figure 12f and gives indications of the subtle interplay between largely passive subthreshold interaction and active interaction via back-propagating action potentials between the soma and apical tuft. At low basal amplitudes (below around 0.3 nA), the probability of an initial somatic spike increases rapidly with subthreshold apical input (below around 1.0 nA) in the short cell, resulting in an increase in burst probability at apical amplitudes below 1.0 nA, but not in the long cell, even though the apical threshold for BAC-firing is higher in the short cell. Once the basal amplitude is sufficient to start reliably generating somatic spikes in the long cell (above 0.3 nA), burst firing probability increases rapidly with apical amplitude due to the lower BAC-firing threshold in this cell. Thus the burst probability in the long cell quickly overtakes that in the short cell for basal amplitudes above 0.3 nA (Figure 12f). This is augmented by the larger apical contribution to generating a short two-spike burst at higher basal amplitudes in the long cell (above around 0.6 nA).

The passive and active differences between the short and long cells are further detailed by the data in Table 1 in the Appendix.

### PID analysis

The change in burst probability landscape with trunk length is also reflected in the PID analysis of information transmission by bursts. Figure 13 shows the PID surfaces for the cell with the short trunk length. Comparing this with Figure 3, it is apparent that each component of information has a mirror-image landscape between long and short trunk lengths.

**Figure 13.**
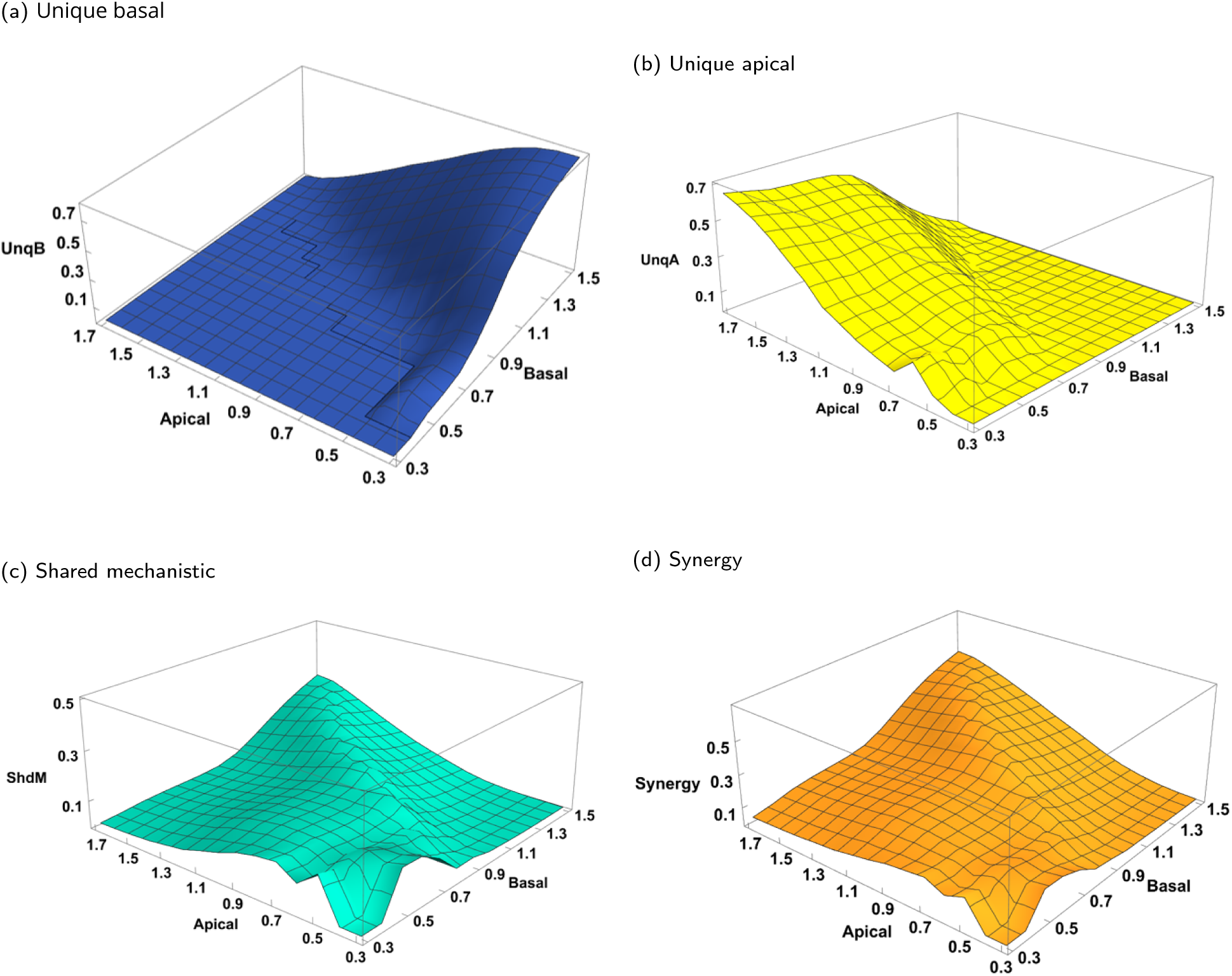
Normalised information-carrying PID quantities for cell with a short trunk length of 200*µ*m, obtained by using the Ibroja method for pairs of basal and apical amplitude ranges: basal ranges from (0, 0.3), (0, 0.4) to (0, 1.5) nA; apical ranges from (0, 0.3), (0, 0.4) to (0, 1.7) nA, both in increments of 0.1 nA. (a) Unique basal. (b) Unique apical. (c) Shared mechanistic. (d) Synergistic.

It appears that with a short trunk the modulatory regime has switched from HL to LH. That is, now it is a low basal input that modulates the burst probability over the full range of apical inputs. Compare particularly Figure 13a,b with Figure 3a,b. In the short cell, little or no unique basal information is transmitted by basal amplitudes below around 0.9 nA, whereas the unique apical information is non-zero and rather constant across this basal range, and rises with apical amplitude. This is due basal input acting to modulate the initial somatic spike probability due to apical input, which then leads to BAC-firing as a function of apical amplitude. At higher basal amplitudes, low apical input serves to modulate two-spike burst probability due to basal input, leading to the rise in unique basal information at basal amplitudes above 0.9 nA.

## Discussion

Using computational models of neocortical layer 5 pyramidal cells (***Bahl et al., 2012***; ***Hay et al., 2011***) that can exhibit output spike bursting via BAC-firing, we have shown how spatially-targetted inhibition and neuromodulation of membrane-bound ion channels can influence the signal interaction of the two prominent excitatory input streams to the basal and apical dendrites, respectively, and the subsequent generation of a burst of output spikes. We have shown previously (***Graham et al., 2025***) that the mode of burst generation depends on the relative strengths of the two excitatory input streams and that several different operating regions can be defined by restricting the amplitude ranges of one or other of the streams. Here we demonstrate that these regions can emerge through the application of inhibition or ion channel modulation, without any change in the amplitude ranges of the excitatory inputs.

### Inhibitory control of two-stream signal interaction

We have demonstrated that spatially-targetted inhibition to the apical tuft and basal dendrites significantly alters the information transmitted in the output bursting probability about the two excitatory input streams, largely independently. Moderate apical inhibition reduces the excitatory apical input to essentially modulating the information transmitted about the basal input, and vice-versa for basal inhibition. Strong inhibition will silence excitatory activity in either area altogether. ***Leleo and Segev (2021)*** have carried out a detailed modelling study of the control of bursting by spatially-distributed inhibition in a morphologically-complex layer 5 pyramidal cell model. The sources of such inhibition and their activation in cortical circuits during cognition is the subject of extensive ongoing study (***Gentet, 2012***; ***Kubota et al., 2016***; ***Wang and Yang, 2018***). ***Phillips (2023)*** gives a review in the light of contextually-modulated information processing and we discuss key points here. Basal inhibition is provided by parvalbumin positive (PV) interneurons that are driven by feed-forward thalamo-cortical input carrying information about sensory stimuli from the external world. Apical inhibition is more diverse and includes somatostatin positive (SST) and neurogliaform (NGF) cells driven by higher cortical regions carrying information about the internal world, encompassing learnt experience and knowledge. Further, pyramidal cells may be disinhibited by vasoactive intestinal peptide positive (VIP) interneurons that selectively inhibit SST interneurons.

Untangling the connectivity and activity patterns of these interneurons within cortical microcircuits is the subject of both experimental and theoretical work (***Jiang et al., 2013***; ***Keller et al., 2020***; ***Shen et al., 2022***; ***Wang and Yang, 2018***). In particular, the role of disinhibition in gating contextual influence from higher cortical areas has been shown to underpin contextually-modulated responses to visual stimuli (***Keller et al., 2020***) and is proposed as a general principle allowing multi-pathway gating of information between brain areas (***Wang and Yang, 2018***). Disinhibition is focal within a cortical column (***Jiang et al., 2013***) and is source-specific, with cross-modal contextual inputs preferentially disinhibiting their target pyramidal cells, whereas within the same modality such inputs largely inhibit their target PCs (***Shen et al., 2022***).

### Neuromodulatory control of signal interaction

Of the myriad effects of neuromodulation, we have investigated the regulatory effects on membrane-bound calcium channels and HCN channels.

### Calcium channels

The dendritic calcium spike is fundamental to the apical contribution to output burst firing. Down-regulation of calcium channels, modelled here as a decrease in their maximum conductance, results in a reduction in influence of excitatory apical input on bursting. A similar reduction in high-frequency firing in response to a volley of synaptic input is seen in the model of ***Shai et al. (2015)***. Apical input now has a largely modulatory influence on bursting generated by basal excitatory inputs. The bursting profile is very similar to that achieved by moderate apical inhibition.

Activation of muscarinic acetylcholine receptors in the apical tuft of layer 5 pyramidal cells promotes calcium spiking through upregulation of R-type calcium channels (***Williams and Fletcher, 2019***). Acetylcholine levels are high during wakefulness and thus could contribute apical-calcium-spike-mediated contextually-modulated information processing in a pyramidal cell (***Aru et al., 2020***). Acetylcholine levels are even higher during REM sleep, potentially leading to apical-only output bursting which could underly dreaming (***Aru et al., 2020***).

### HCN channels

The effect of HCN channel activation and its modulation on output bursting is much more complex. Our modelling shows that with brief excitatory stimuli arriving during resting conditions in the pyramidal cell, HCN channels with high density in the apical dendrites may either amplify or reduce the probability of bursting as a function of the strength of apical input. In the ***Bahl et al. (2012)*** cell model the dendritic calcium spike is mediated by high-voltage-activated (HVA) calcium channels alone, and the HCN channels amplify the effect of apical input by depolarising the apical tuft at rest. However, addition of a tonic leak current due to a conductance that is proportional to the maxiumum H channel conductance can turn the effect of the HCN channels into one of inhibition (***Migliore and Migliore, 2012***). Alternatively, in the ***Hay et al. (2011)*** model, low-voltage-activated (LVA) channels contribute significantly to the calcium spike, and the tuft depolarisation due to the HCN channel activity now inactivates these channels and reduces the ability of apical input to generate a calcium spike. This also has been investigated extensively in the ***Hay et al. (2011)*** model by ***Mäki-Marttunen and Mäki-Marttunen (2022)***.

Experimental data indicates that HCN channels do have an inhibitory effect on dendritic stimulation from rest (***Harnett et al., 2015***; ***Labarrera et al., 2018***) and that this is likely due to the shunting effect of the resting HCN channel conductance in the dendrites, without LVA calcium channel inactivation playing a role. They also show that HCN channel activation actually amplifies the effect of perisomatic excitatory stimulation, where the depolarisation due to HCN channel activation is more significant than the current shunt, with also a contribution from passive current flow from the depolarised dendrites ***Harnett et al. (2015)***. This is seen with the ***Hay et al. (2011)*** model in the simulations of ***Mäki-Marttunen and Mäki-Marttunen (2022)*** and in our results from both the ***Bahl et al. (2012)*** and Hay et al. (2011) models, where bursting by strong basal input is inhibited with the removal of the HCN channels (Figures 10 and 11).

However, in the dynamic situation of time-varying membrane potential, dendritic hyperpolarisation preceding brief excitatory input results in HCN channels amplifying calcium spiking through rebound excitation. This occurs even if the calcium spike is generated through LVA channels, as a preceding hyperpolarisation also acts to deinactivate these channels. Thus neuromodulatory regulation of the HCN channels could either amplify or suppress bursting through dendritic calcium spiking, depending on the membrane physiology and the patterns of ongoing synaptic activity. Due to active conductances, the apical dendrites respond preferentially to specific frequencies of synaptic input (***Kalmbach et al., 2017***). In particular, the Ih current generated by HCN channels can contribute to theta-frequency (4-10 Hz) resonance in pyramidal cell dendrites (***Labarrera et al., 2018***). Application of an adrenergic receptor agonist in the somatosensory cortex of awake mice increases tuft excitability and the likelihood of calcium spiking, putatively through modulation of dendritic HCN channels (***Labarrera et al., 2018***). Noradrenaline levels are high during wakefulness and thus, like acetylcholine, this may contribute to contextually-modulated information processing (***Aru et al., 2020***).

In addition to affecting the generation of calcium spikes, neuromodulation may also influence the electrical coupling between the apical tuft and cell body and thus the effect of a distal calcium spike on output bursting. Thalamic activation of metabotropic glutamate receptors and activation of muscarinic acetylcholine receptors along the apical dendritic trunk promotes signal transfer from the tuft to the soma (***Suzuki and Larkum, 2020***). Neurogliaform interneurons activate dendritic GABA*_B_* receptors that in turn activate dendritic potassium channels (GIRK) and thus lead to a reduction in signal transfer from the tuft to the soma (***Schulz et al., 2021***).

### Effect of apical trunk length on signal interaction

Reducing the trunk length in the ***Bahl et al. (2012)*** while maintaining the electrophysiological gradients of ion channels highlights the subtle interplay between active and passive (subthreshold) signal exchange between the apical tuft and soma. In the shorter cell, the back-propagating sodium action potential (bAP) is actually less potent in facilitating BAC-firing. This has been shown by ***Galloni et al. (2020)*** to be due to broadening of the active bAP as it travels to the tuft resulting in greater charge transfer to the tuft in a longer cell. So there is a balance between the threshold for BAC-firing and the potentcy of a calcium spike in the tuft being able to stimulate multiple somatic sodium spikes that is a function of trunk length.

Experiments in thick-tufted layer 5 pyramidal cells show that BAC-firing and subsequent bursting is limited to the largest cells (***Fletcher and Williams, 2019***; ***Galloni et al., 2020***). There is a rostral-caudal gradient from large to small cells and their bursting characteristics in primary visual cortex (***Fletcher and Williams, 2019***) and a similar difference between primary (larger cells) and secondary (smaller cells) visual areas (***Galloni et al., 2020***) in the rat. At the other extreme, human pyramidal cells are larger and consequently could exhibit even greater compartmentalisation of the distal apical dendrites. Human thick-tufted layer 5 pyramidal cells can have similar morphological and electrophysiological characteristics to rodent cells and do show dendritic plateau potentials (***Kalmbach et al., 2021***) but it is not clear if these may lead to bursting in the soma (***Beaulieu-Laroche et al., 2018***).

### Conclusion

Two-pathway signal integration and subsequent burst firing in layer 5 pyramidal cells involves active processes in the perisomatic (basal) and apical tuft dendrites and the active and passive interaction between these compartments through the apical dendritic trunk. The burst firing landscape as a function of the strength of basal and apical excitatory inputs may be shaped by spatially-targetted inhibition and neuromodulation of ion channel activity in each of these three compartments (perisomatic, tuft and trunk dendrites), resulting in distinct information transmission characteristics for the cell.

## Appendix

### Passive and active cell properties

Table 1 details the amplitudes of passive signal propagation between the soma and apical tuft and the thresholds for initiating somatic sodium and apical tuft calcium spikes, for the original cell with a 500 *µ*m apical trunk length and for a shorter cell with a 200 *µ*m trunk length.

As would be expected, the shorter trunk length allows greater subthreshold interaction between the soma and tuft, with stimuli at either site resulting in a larger depolarisation at the other site in the shorter cell (Table 1, rows 1-2). This results in the basal (soma) amplitude threshold for generating single somatic spikes decreasing faster in the short cell as the apical amplitude is increased from zero (Table 1, rows 6-8). Thus, in the noisy cell the single somatic spike probability rises more rapidly with increasing basal and apical amplitudes in the short cell.

However, counter-intuitively, the tuft depolarisation due to a somatic spike is larger in the long cell (Table 1, rows 3-5) due to sodium-channel-dependent active back-propagation. Consequently, the somatic (basal) amplitude required to generate a short, two-spike burst in the soma is lower in the longer cell due to a contribution from apical depolarisation (Table 1, row 9), even though apical-to-soma attenuation is greater in the long cell. Further, following an initial somatic spike, the apical input amplitude required to initiate BAC-firing is increased in the short cell due to the lower tuft depolarisation resulting from the somatic spike (Table 1, row 10). Note that the threshold apical current amplitude alone required to generate a calcium spike in the tuft is unchanged with trunk length ((Table 1, row 11). This relationship between trunk length and and tuft excitability is explored by ***Galloni et al. (2020)*** and explained by them as due to the active effects of dendritic sodium channels on the back-propagating action potential.

### Transfer function definition

The transfer functions for the different bursting regimes are defined as follows. Full details of how these functions are derived are to be found in ***Graham et al. (2025)***.

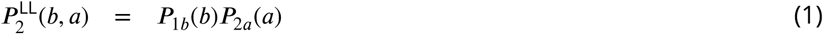

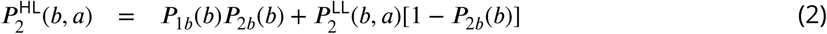

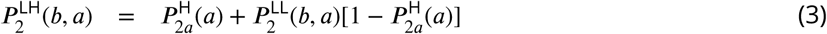

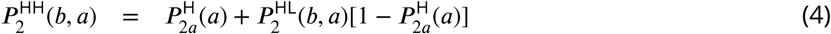

where the individual components are:

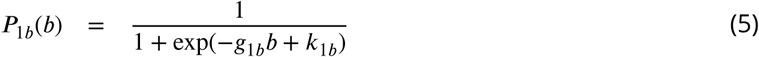

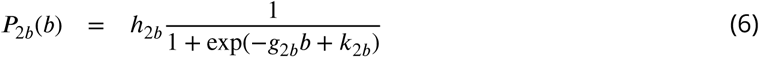

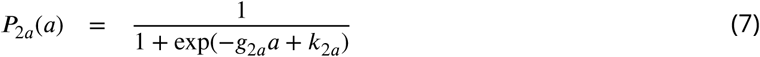

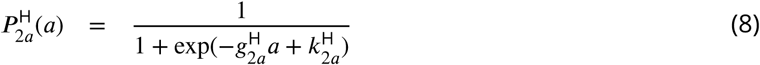

The components can be interpreted as:

*P*_1*b*_(*b*) ≡ *P* (*Z*_1_ = 1l*b*) is the probability of an initial somatic spike (binary random variable *Z*_1_ = 1 when spike occurs) due to basal input alone.

*P*_2*b*_(*b*) ≡ *P* (*Z*_2_ = 1l*b*) is the burst probability (binary random variable *Z*_2_ = 1 when second spike (burst) occurs) due to basal input alone.

*P*_2*a*_(*a*) ≡ *P* (*Z*_2_ = 1l*Z*_1_ = 1*, a*) is the contribution of apical input to a full (or partial) calcium spike which can lead to a second (or more) somatic spike, following an initial somatic spike.

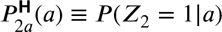 is the probability that apical input produces a calcium spike on its own.

### Transfer function fitting

Table 2 contains the fitted parameter values for the different cases. The full HH transfer function is fit in all cases, and reduced transfer functions HL, LH and LL are fit where appropriate.

For comparison with the cases where subregimes HL, LH and LL are approximated either through dendritic inhibition or down-regulation of calcium channels, Figure 14 and Figure 15 show the fits to these cases using the full HH transfer function. These are discussed in the main text and can be compared with Figure 4 and Figure 8.

**Figure 14.**
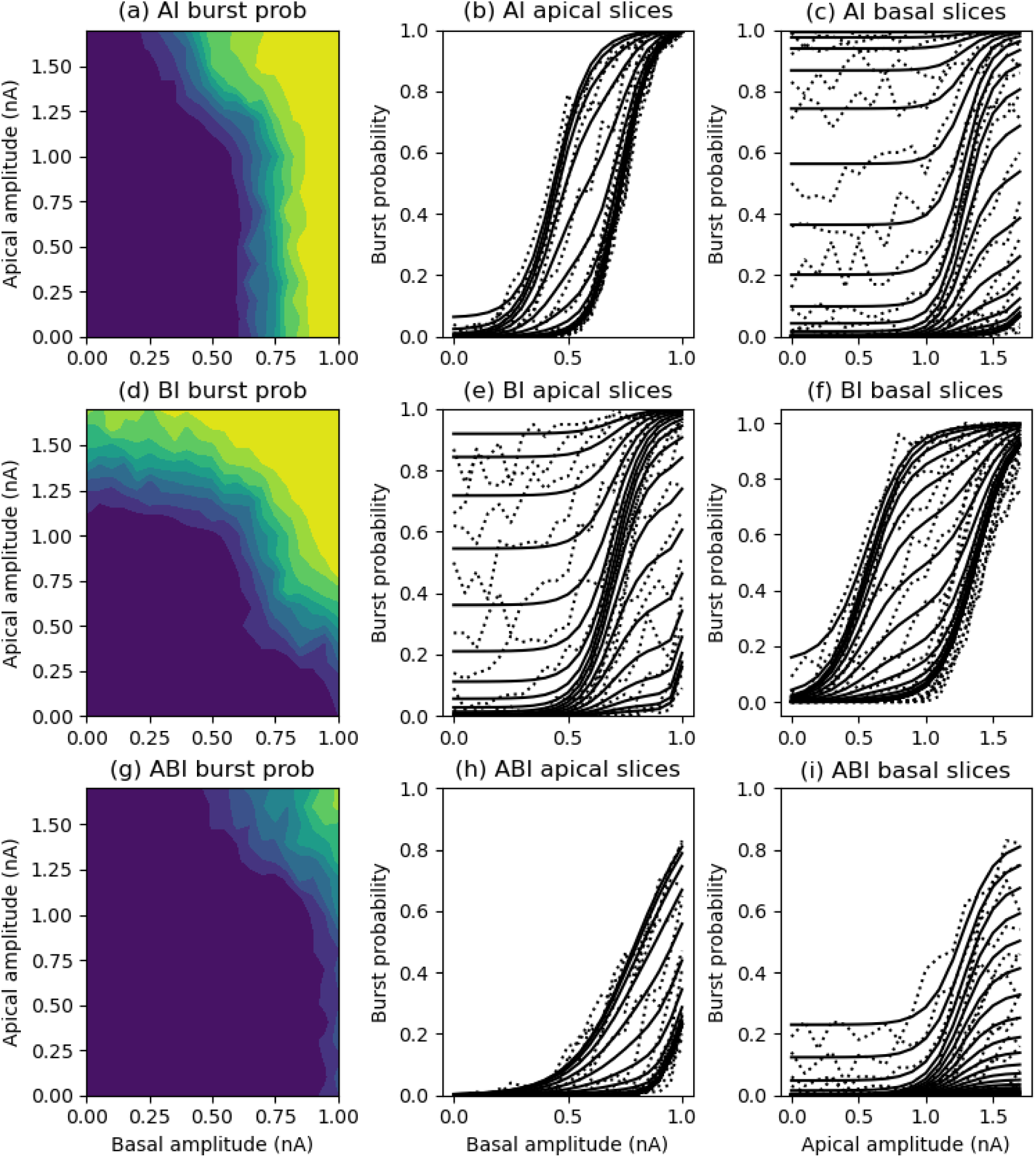
Contour and slice plots of burst probability from the simulation data for the HH amplitude ranges but with dendritic inhibition. (a) Tonic inhibition of 0.01 *µS* applied to the apical tuft. (b) Tonic inhibition of 0.01 *µS* applied to the basal dendrites, with basal excitation shifted to the basal dendrites. (c) Tonic inhibition of a strength of 0.007 *µS* applied to both the basal dendrites and apical tuft. Solid lines in the slice plots are transfer function fits using the full HH transfer function in each case.

**Figure 15.**
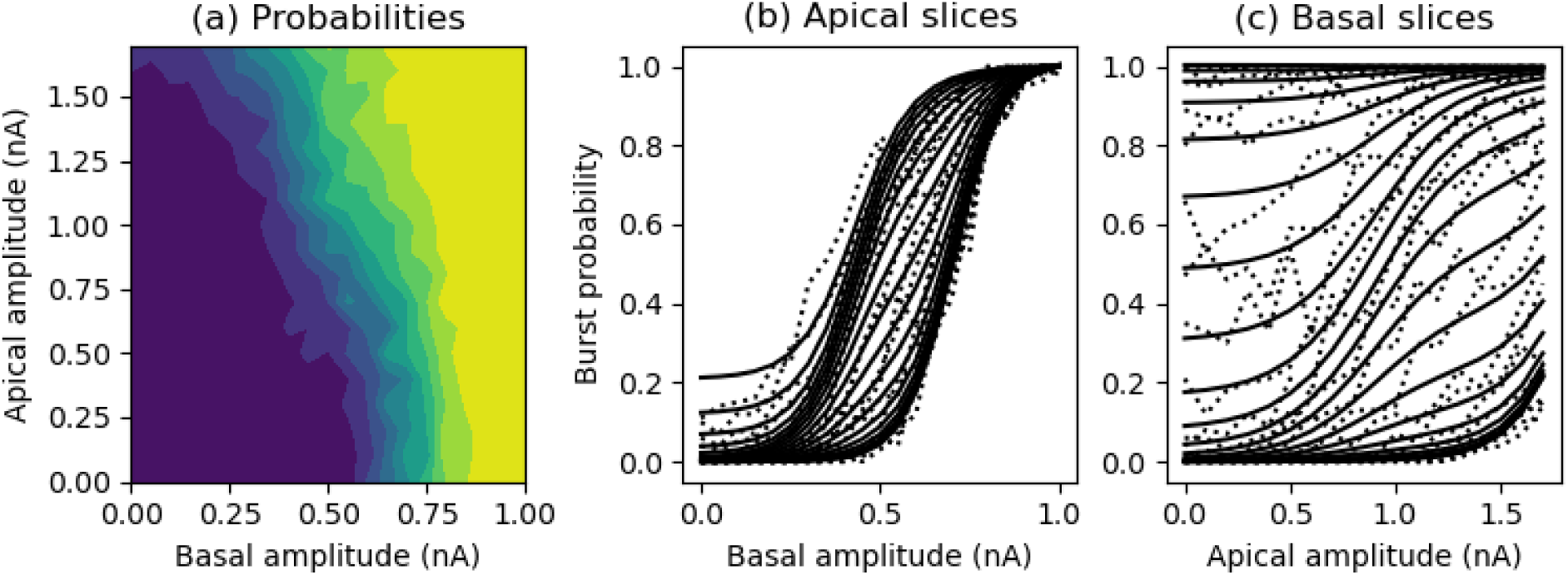
(a) Contour plots of burst probability from the simulation data for the HH amplitude ranges with tuft calcium channel density reduced to one quarter of its original value. (b) Simulation (dotted lines) and fitted HH transfer function (solid lines) curves for burst probability as a function of basal amplitude for individual values of apical amplitude. (c) As for (b) but as a function of apical amplitude for values of basal amplitude.

### PID analysis over fixed ranges

To give a further comparison of the effects of inhibition and neuromodulation on information transfer as estimated using partial information decomposition (PID), Figure 16 shows the PID over the full range of apical and basal amplitudes used in the different situations. These PIDs are shown alongside those for the comparable PIDs from the original simulation data over suitably reduced ranges. No attempt was made to optimise the strength of inhibition or neuromodulation used to match the original PIDs, but the comparability is striking, with qualitative similarity across each PID component. The main quantitative difference is that the inhibition applied to the basal dendrites alone does not produce as much unique information about the apical input, but does show the same trend as the LH regime.

**Figure 16.**
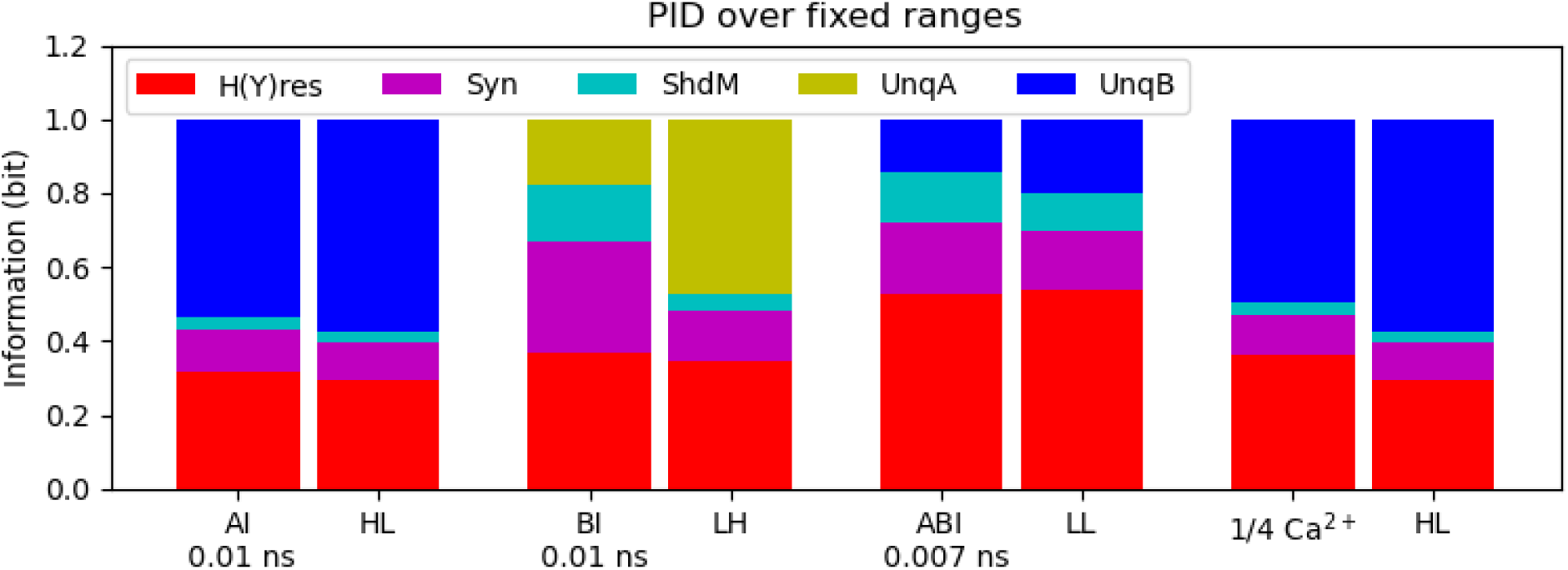
PID over the full amplitude ranges when inhibition is applied to the apical dendrites (AI), the basal dendrites (BI), both dendrites (ABI), or with a reduction in the tuft calcium channel density (1/4 Ca^2+^), compared with the PID over restricted amplitude ranges when no inhibition or calcium channel reduction is applied (HL, LH, LL, HL). H(Y)res is residual entropy; Syn is synergistic; ShdM is shared mechanistic; UnqA is unique apical and UnqB is unique basal information.

